# Termination factor Rho mediates transcriptional reprogramming of *Bacillus subtilis* stationary phase

**DOI:** 10.1101/2022.02.09.479721

**Authors:** Vladimir Bidnenko, Pierre Nicolas, Cyprien Guérin, Sandra Dérozier, Arnaud Chastanet, Julien Dairou, Yulia Redko-Hamel, Matthieu Jules, Elena Bidnenko

**Affiliations:** Micalis Institute, INRAE, AgroParisTech, Université Paris-Saclay, 78350 Jouy-en-Josas, France; MaIAGE, INRAE, Université Paris-Saclay, 78350 Jouy-en-Josas, France; Laboratoire de Chimie et Biochimie Pharmacologiques et Toxicologiques, CNRS, UMR 8601, Université de Paris, F-75006, Paris, France.

## Abstract

Reprogramming of gene expression during transition from exponential growth to stationary phase is crucial for bacterial survival. In the model Gram-positive bacterium *Bacillus subtilis*, this process is mainly governed by the activity of the global transcription regulators AbrB, CodY and Spo0A. We recently showed that the transcription termination factor Rho, known for its ubiquitous role in the inhibition of antisense transcription, is involved in Spo0A-mediated regulation of differentiation programs specific to the stationary phase in *B. subtilis*. To identify other aspects of the regulatory role of Rho during adaptation to starvation, we have constructed a *B. subtilis* strain that expresses *rho* at a relatively stable high level in order to circumvent its decrease occurring in the wild-type cells entering the stationary phase. We show that *B. subtilis* cells stably expressing Rho fail to sporulate and to develop genetic competence, which is largely, but not exclusively, due to abnormally low expression of the master regulator Spo0A. Moreover, in addition to a global decrease of antisense transcription, these cells exhibit genome-wide alterations of sense transcription. A significant part of these alterations affects genes from global regulatory networks of cellular adaptation to the stationary phase and reflects the attenuated de-repression of the AbrB and CodY regulons and the weakened stringent response. Accordingly, stabilization of Rho level reprograms stationary phase-specific physiology of *B. subtilis* cells, negatively affects cellular adaptation to nutrient limitations and alters cell-fate decision-making to such an extent that it blocks development of genetic competence and sporulation. Taken together, these results indicate that the activity of termination factor Rho constitutes a previously unknown layer of control over the stationary phase and post-exponential adaptive strategies in *B. subtilis*, from the adjustment of cellular metabolism to the activation of survival programs.

## Introduction

Transcription termination is a critical step in regulation of gene expression in all living organisms. In bacteria, termination is achieved by two mechanisms: factor-independent, which is associated with specific sequence forming an RNA terminator hairpin, and factor-dependent, which relies mostly on the activity of an RNA helicase–translocase, transcription termination factor Rho [1–3].

Since its initial characterization in 1993 [4], termination factor Rho was repeatedly shown to be dispensable for the model Gram-positive bacterium *Bacillus subtilis* in laboratory growth conditions [5–8]. On the contrary, the viability of numerous Gram-negative bacteria depends strictly on active Rho (reviewed in [9]). While, the essentiality of the *rho* gene varies among different bacterial species, Rho is recognized now as the major factor controlling pervasive antisense transcription in both Gram-positive and Gram-negative bacteria [6, 10–12]. Moreover, Rho inactivation alters significantly the expression of protein-coding genes by a combination of direct *cis* and indirect *trans* effects in bacterial species in which the *rho* gene is non-essential, as reported for *B. subtilis*, *Staphylococcus aureus*, and *Bacillus thuringiensis* [8, 11, 12].

In *B. subtilis*, we have shown, using *Δrho* mutant, that a significant part of the Rho-controlled transcription is connected to the regulation of three mutually exclusive differentiation programs: cell motility, biofilm formation, and sporulation. To a large extent, the choice between these and other cell fates (e.g., genetic competence, cannibalism toxins production) upon entry into the stationary phase depends on cellular levels of the phosphorylated active form of the master regulator Spo0A (Spo0A∼P). Only cells expressing high levels of Spo0A∼P can commit to sporulation, an ultimate survival option of *B. subtilis* cells at stationary phase [14, 15]. We have established that deletion of the *rho* gene prevents Rho-dependent intragenic termination of the *kinB* transcript encoding the sensory kinase KinB, thereby activating the Spo0A phosphorelay and increasing cellular levels of Spo0A∼P to a threshold triggering sporulation. Thus, Rho inactivation increases the efficiency of sporulation and inhibits the alternative cell fates [8].

In the human pathogens *S. aureus, Mycobacterium tuberculosis* and *Clostridioides difficile*, Rho inactivation induces the expression of virulence factors essential for the successful host colonization and infection [12, 16, 17]. Likewise, Rho affects expression of genes involved in cellular differentiation, colonization and pathogenesis in *B. thuringiensis* [13]. Overall, these data indicate that in *B. subtilis* and other Gram-positive bacteria, Rho plays an important role in the regulation of different phenomena associated with the stationary phase.

This specific physiological state of growth caused by nutrients depletion is characterized by slowdown of macromolecular synthesis, reorientation of the cellular metabolism towards alternative metabolic pathways, activation of the stringent response and alternative sigma factors [18].

Along with Spo0A, two other key transcriptional regulators, AbrB and CodY, sensing environmental and intracellular metabolic status drive the reprogramming of metabolism and the initiation of stationary phase-specific developmental programs in *B. subtilis* [19]. During exponential growth, AbrB suppresses transcription of over two hundred genes that are switched ON upon AbrB depletion during the transition to the stationary phase [20–22]. AbrB plays an important role in the interconnected regulatory networks governing the initiation of sporulation and development of genetic competence by controlling the expression of transition-phase sigma factor SigH and competence transcription factor ComK [23, 24]. Thus, AbrB depletion is important for cells entering the stationary phase. Expression of the *abrB* gene is repressed by Spo0A∼P, which also indirectly controls AbrB DNA binding activity [21, 25, 26]. In addition, AbrB is an unstable protein and its concentration decreases rapidly due to degradation of the *abrB* mRNA triggered by small regulatory RNA, RnaC [27].

The pleiotropic regulator CodY directly and indirectly represses transcription of the numerous genes required for adaptation to nutrient limitation [28, 29]. This repression is released to activate the alternative nutrient acquisition pathways when cells enter into the stationary phase [30, 31]. CodY is also implicated in the control of genetic competence and sporulation [32, 33]. CodY modulates its own DNA-binding affinity by sensing two metabolites: branched-chain amino acids (BCAA) isoleucine, leucine, and valine and the nucleoside triphosphate GTP. In the absence of any of these ligands, the ability of CodY to bind DNA is impaired [30, 34]. In that way, activity of CodY is linked to stringent response, a widespread stress resistance mechanism essential for stationary phase survival [35–37]. It is characterized by the synthesis of the alarmone guanosine-(penta)tetra-phosphate ((p)ppGpp), mainly provided by a bifunctional synthetase/hydrolase Rel sensing starved ribosomes [38–40]. In *B. subtilis*, (p)ppGpp modifies genome-wide transcription indirectly, by causing a decrease in GTP levels due to the inhibition of activity of GTP-synthesizing enzymes and consumption of GTP during synthesis of (p)ppGpp [41–43]. A decrease in the GTP levels causes de-repression of genes from the CodY regulon, negatively affects transcription from promoters of stable RNA synthesis genes (e. g., genes involved in the ribosome biogenesis) and re-directs RNA polymerase from these GTP-initiating promoters to promoters of biosynthetic genes [44–46]. In addition, (p)ppGpp directly represses the activity of the DNA primase, thereby regulating DNA replication [47] and inhibits protein synthesis [48–50]. Furthermore, it is known that burst of (p)ppGpp upon entering the stationary phase contributes to the induction of genetic competence and sporulation [33, 51, 52].

It is important to note that in the wild type cells *rho* mRNA and Rho protein levels decrease during transitional and stationary growth phases [6, 8, 11]. By analogy with AbrB, this phase-dependent depletion of Rho suggests that, in addition to controlling long-term survival strategies, Rho may participate in the regulation of the earlier stages of physiological adaptation to stationary phase and, in a broader sense, in the control of transition and stationary phase-specific transcriptomes. Such hypothetical Rho activity could not be detected using the Δ*rho* mutant. Thus, we assumed that the stabilization of Rho levels over an extended period of growth would provide an experimental model to study this aspect of Rho regulatory activity and thereby to expand our knowledge about Rho-mediated regulation of gene expression and its effect on the cellular physiology of *B. subtilis*.

To evaluate this hypothesis, we used a combination of experimental approaches, including genome-wide transcriptional analysis, monitoring of time-course activity of selected promoters, morphological and functional studies of the *B. subtilis* strain (hereinafter, Rho^+^), in which Rho was maintained at a relatively stable high level throughout exponential and stationary growth phases. We show that stable expression of the *rho* gene causes system-level alterations of genome transcription. Rho reprograms cellular physiology and starvation-specific developmental programs driven by the activity of key transcriptional regulators AbrB, CodY, ComK, and Spo0A. Moreover, Rho^+^ strain exhibits partially relaxed phenotype characterized by a weakened stringent response.

Altogether, these findings provide new functional insights into the role of the transcription termination factor Rho in the physiology of *B. subtilis* and indicate that strict regulation of Rho expression is crucial for the functionality of the complex gene networks governing the stationary-phase adaptation in this model Gram-positive bacterium.

## Results

### Heterogenic expression system assures Rho expression at a steady high level

To evaluate the impact of the stabilization of Rho levels on stationary phase-specific phenomena, we first conceived a system that would maintain relatively stable Rho amounts over exponential and stationary growth. The plasmid that we previously used for Rho over-expression seemed unsuitable for this purpose due to the copy number heterogeneity [8, 53], and the presence of the 5’UTR region of *rho* gene previously implied in auto-regulation of Rho [5]. To overcome these limitations, we have disconnected the expression of *rho* from any regulatory circuits acting at its natural locus by placing a copy of the *rho* gene under the control of heterogenic expression signals at an ectopic location. Briefly, we substituted the native *rho* promoter with a well-characterized constitutive *B. subtilis* promoter P*_veg_* [54, 55], replaced the 5’UTR of the *rho* gene with the ribosome binding site of the *tagD* gene [56] and placed this *rho* expression unit at the *amyE* locus. Expression of the *rho* gene driven by these regulatory elements had no effect on growth rate of the resulting *B. subtilis* strain (hereinafter, Rho^+^) in rich LB medium over an extended period (∼10 h), ranging from exponential into the stationary phase (Fig 1A).

**Fig 1.**
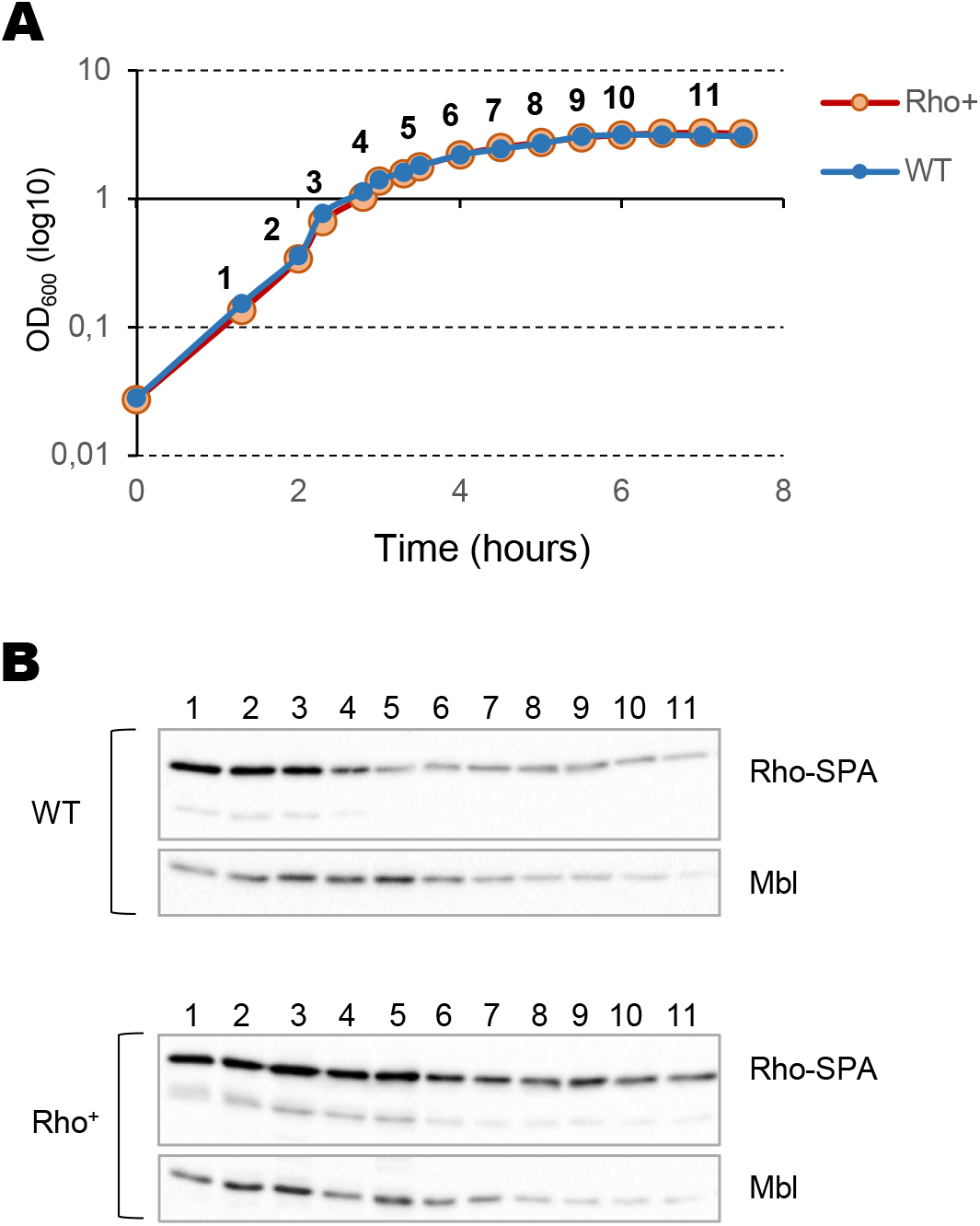
Heterogenic expression of the transcription termination factor Rho. *B. subtilis* cells expressing SPA-tagged Rho protein from natural (WT) and Pveg (Rho^+^) promoters were grown in LB medium (**A**) and analyzed for Rho protein content at the indicated time points by immunoblotting with ANTI-FLAG M2 monoclonal antibodies (Rho-SPA; **B**). Equal amounts of total protein extracts were loaded onto the gel according to Bradford assay; samples’ equilibrium between the strains at each time-point and the quality of transfer were controlled by visualization of Mbl protein using specific anti-Mbl antibodies (Mbl; **B**). Note that the Mbl levels decrease in the stationary phase.

To evaluate expression levels of Rho protein under these growth conditions, we used the relative to WT and Rho^+^ strains expressing the SPA-tagged Rho protein to reveal Rho content by immunoblotting [8]. In the WT cells, Rho-SPA protein levels steadily decreased during transition into the stationary phase in LB medium (Fig 1B). In contrast, Rho^+^ cells showed relatively stable levels of Rho-SPA during the exponential growth and after entering the stationary phase (Fig 1B). The decrease of Rho-SPA observed in the Rho^+^ cells during the late stationary phase could be due to a decline of the P*_veg_* promoter’s activity at this stage as reported previously [55].

We concluded that using the heterogenic expression system assures a steady high level of Rho expression in the stationary-phase *B. subtilis* cells.

### Rho^+^ strain exhibits sporulation-deficient phenotype

We initiated the analysis of physiological effects of a steady high Rho content by assessment of the sporulation capacity of Rho^+^ cells.

The WT, Δ*rho* mutant (RM) and Rho^+^ cells were cultured in the sporulation-inducing DS medium and compared for the ability to form heat-resistant spores (Materials and Methods). Depending on the experiment, 20% to 40% of the WT cells developed spores under the used conditions, while *rho* deletion increased the sporulation rate to its maximum as previously reported (Fig 2A; [8]). In contrast, the efficiency of spore formation by the Rho^+^ strain was reduced up to 10^-5^. The rare spores isolated from Rho^+^ cultures appeared to be suppressor mutants able to form thermo-resistant spores with a near-WT efficiency (S1 Table).

**Fig 2.**
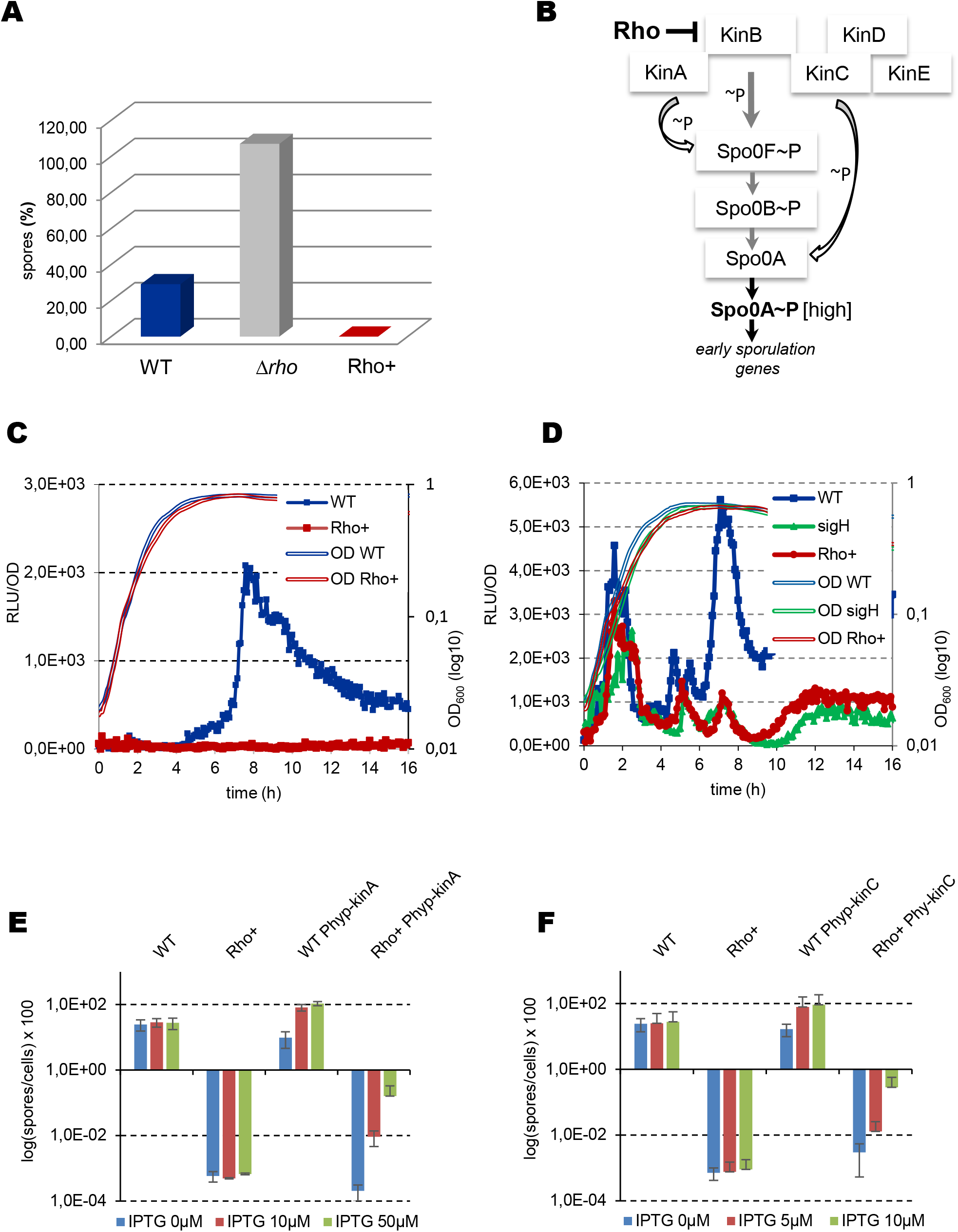
*B. subtilis* Rho^+^ strain exhibits sporulation-negative phenotype. (**A**) Sporulation efficiency of *B. subtilis* WT, Δ*rho* and Rho^+^ cells. Cells inoculated at OD_600_ 0.025 were grown in DS medium at 37° for 20 hours and analyzed for heat-resistant spores as described in Materials and Methods. Sporulation efficiency was estimated as proportion of viable cells in the heated and unheated cultures. Plotted are the mean values from four independent experiments with three biological replicas for each strain, and the SDs ≤ 10%. (**B**) Schematics of the multicomponent Spo0A phosphorelay. Only the key elements relevant to this study are shown. Phosphoryl groups are transferred from sensor protein kinases (KinA-E) to Spo0F, Spo0B, and ultimately to Spo0A. Sporulation is triggered when the level of Spo0A∼P reaches a high threshold level. The bar-headed line indicates negative Rho-mediated regulation of KinB expression. (**C**) Kinetics of luciferase Luc expression from the promoter of early sporulation gene *spoIIAA* in *B. subtilis* WT (blue line) and Rho^+^ (red line) cells induced for sporulation. (**D**) Kinetics of luciferase Luc expression from the promoters of *spo0A* gene in WT (blue line), Rho^+^ (red line) and *sigH* mutant (green line) cells induced for sporulation. In **C** and **D**, cells bearing transcriptional fusions *spoIIAA-luc* and *spo0A-luc*, respectively, were grown in DS medium and analyzed for luciferase activity at five-minutes intervals in a multimode microplate reader as described in Materials and Methods. For each strain, plotted are the mean values of luminescence readings corrected for OD_600_ from four independent cultures analyzed simultaneously (solid lines with symbols) and characteristic growth curves (double-lined) measured by OD_600_. The experiments were reproduced at least three times. The results from the representative experiments are presented. (**E** and **F**) Synthetic over-production of sensor histidine kinases KinA or KinC does not rescue sporulation-negative phenotype of Rho^+^ cells. Sporulation efficiency of *B. subtilis* WT and Rho^+^ strains and their respective derivatives expressing *kinA* (**E**) and *kinC* (**F**) genes under control of the IPTG-inducible promoter. Cells were inoculated at OD_600_ 0.025 in DS medium containing IPTG at the indicated concentrations and grown at 37° during 20 hours; sporulation efficiency was analyzed as described above (Fig 2A) and in Materials and Methods. Plotted are the mean values from three independent experiments with three biological replicas of each strain (rectangulars) with standard deviation SD (bar-headed lines).

Analysis of eight independent sporulation-proficient Rho^+^ suppressors revealed six different point mutations within the *rho* coding sequence of the ectopic *rho* expression unit; two mutations were isolated twice. This reaffirms the determining role of Rho in the sporulation-deficient phenotype of Rho^+^ cells (S1 Table). Two mutant Rho proteins were truncated by a stop codon at the position 146 (Q146Stop) and six others had single amino acid changes: A177T, N274H (isolated twice), G286R, G287R and P335R (S1 Fig). The primary sequence of *B. subtilis* Rho subunit displays some characteristic motifs identified previously by studies of different Rho proteins [57–60]; (S1 Fig). In accordance with these data, three of the identified point mutations might have drastic effect on Rho activity. Replacement of glycine by arginine at the positions 286 and 287 (G286R and G287R, respectively) could destroy the highly conserved Q-loop forming a secondary RNA binding site, while the substitution of alanine177 localized within one of the Walker motifs by threonine (A177T) could affect ATP binding. Indeed, in a complementation assay using the Δ*rho* mutant we showed that suppressor mutations G287R, G286R and A177T completely inactivate Rho protein, while N274H and P335R mutant proteins remain partially active (S1 Table).

Considering that Rho affects sporulation by controlling activity of the Spo0A phosphorelay (Fig 2B); [8], strong inhibition of sporulation in Rho^+^ cells suggested that stabilization of the Rho level during stationary phase effectively suppresses the accumulation of active Spo0A∼P. To assess the activity of Spo0A∼P in Rho^+^ cells, we first analyzed the real-time expression of the *spoIIAA-AB-sigF* operon using the transcriptional fusion of its promoter to the firefly luciferase gene *luc* (P*_spoIIAA_-luc*) [8]. Expression of the *spoIIAA-AB-sigF* operon depends on the alternative sigma factor SigH and is activated at a high threshold level of Spo0A∼P [26, 61–63]. As shown in Fig 2C, whereas in WT cells grown in DS medium the P*_spoIIAA_* promoter was switched ON roughly three hours after the entry into stationary phase, no expression could be detected in Rho^+^ cells, suggesting that cells failed to accumulate sufficient amount of Spo0A∼P.

To further characterize activation of Spo0A in Rho^+^ cells, we analyzed the expression of the *spo0A* gene itself. Transcription of *spo0A* is driven by two promoters, the vegetative SigA-dependent Pv and the SigH/Spo0A-controlled Ps, which is activated at the onset of sporulation at a low level of Spo0A∼P [13, 62, 64, 65]. The reporter P*_spo0A_*-*luc* fusion, which we used for analysis, is established at the natural *spo0A* locus and monitors activity of both Pv and Ps promoters [8, 52].

During exponential growth, the P*_spo0A_*-*luc* expression was rather similar in WT and Rho^+^ cells suggesting that Pv promoter was not affected by Rho (Fig 2D). In contrast, two hours after the entry into stationary phase, P*_spo0A_*-*luc* activity greatly increased in WT cells, but remained low in Rho^+^ cells. We noticed that expression kinetics of the P*_spo0A_-luc* in Rho^+^ cells was very similar to that observed in the *sigH* mutant, in which the activity of Ps promoter is abolished (Fig 2D); [64, 65]. This suggests that promoter Ps of the *spo0A* gene was inactive in Rho^+^ cells either due to the Spo0A∼P level lower than required for promoter’s activation or, not mutually exclusive, due to the low activity of SigH.

Taken together, these results demonstrated that stably expressing *rho* drastically reduces accumulation of the active Spo0A∼P under sporulation-stimulating conditions.

### Synthetic over-production of sensor histidine kinases KinA or KinC does not rescue the sporulation-deficient phenotype of Rho^+^ strain

To investigate whether the sporulation-negative phenotype of Rho^+^ cells was solely due to a low level of active Spo0A∼P, we attempted to boost Spo0A phosphorylation (Fig 2B). To this end, we first used a system over-expressing the major kinase of the phosphorelay, KinA, from an IPTG-inducible promoter (P_hyspanc_-*kinA*); [13]. We transferred this system in WT and Rho^+^ cells and assessed their sporulation in DS medium at different concentrations of IPTG inducer. As shown in Fig 2E, addition of IPTG at 10µM and 50µM (concentrations shown to induce the *kinA* gene to a saturation level [14]), triggered sporulation in ∼100 percent of WT cells. The over-expression of KinA in Rho^+^ cells also increased the sporulation frequency ∼10^3^-fold, which remained, however, much below the sporulation level of the non-induced WT cells (Fig 2E). Thus, artificially triggering the phosphorelay appeared insufficient to restore sporulation in Rho^+^ cells and suggested that other roadblocks could exist either within or outside the phosphorelay.

To relieve the accumulation of Spo0A∼P from the control of phosphorelay, we further used an IPTG-regulated system over-producing the sensor histidine kinase KinC, known to transfer phosphate directly to Spo0A [13, 66, 67]. As shown in Fig 2F, induction of KinC expression at IPTG concentrations of 5 to 10 µM, previously shown to be optimal for proper activation of Spo0A [68], stimulated sporulation in WT cells to maximal levels, but resulted only in a partial restauration of sporulation efficiency in Rho^+^ strain, as in the case of KinA over-production. Overall, we concluded that over-production of sensor kinases ensuring consequent increase of Spo0A phosphorylation either directly (KinC) or via the phosphorelay (KinA) cannot fully suppress the sporulation-negative phenotype of Rho^+^ cells. This indicates that Rho negatively affects sporulation not only by repressing Spo0A activation, but also at other stages.

### Rho^+^ strain exhibits competence-deficient phenotype

The intermediate Spo0A∼P level, which is raised transitionally during the late exponential growth without the activation of the sporulation-specific *spo0A* promoter Ps, was shown to be crucial for development of genetic competence (Fig 3A); [69, 70]. We noticed that, contrarily to WT, Rho^+^ strain was not transformable using a common two-step transformation procedure (Materials and Methods).We set to characterize this phenotype of Rho^+^ cells in more details. Since competence is a transient state, we constantly monitored its development, testing cells for genetic transformation during three hours after they were transferred from a rich defined growth medium to a competence-inducing medium (Materials and Methods).

**Fig 3.**
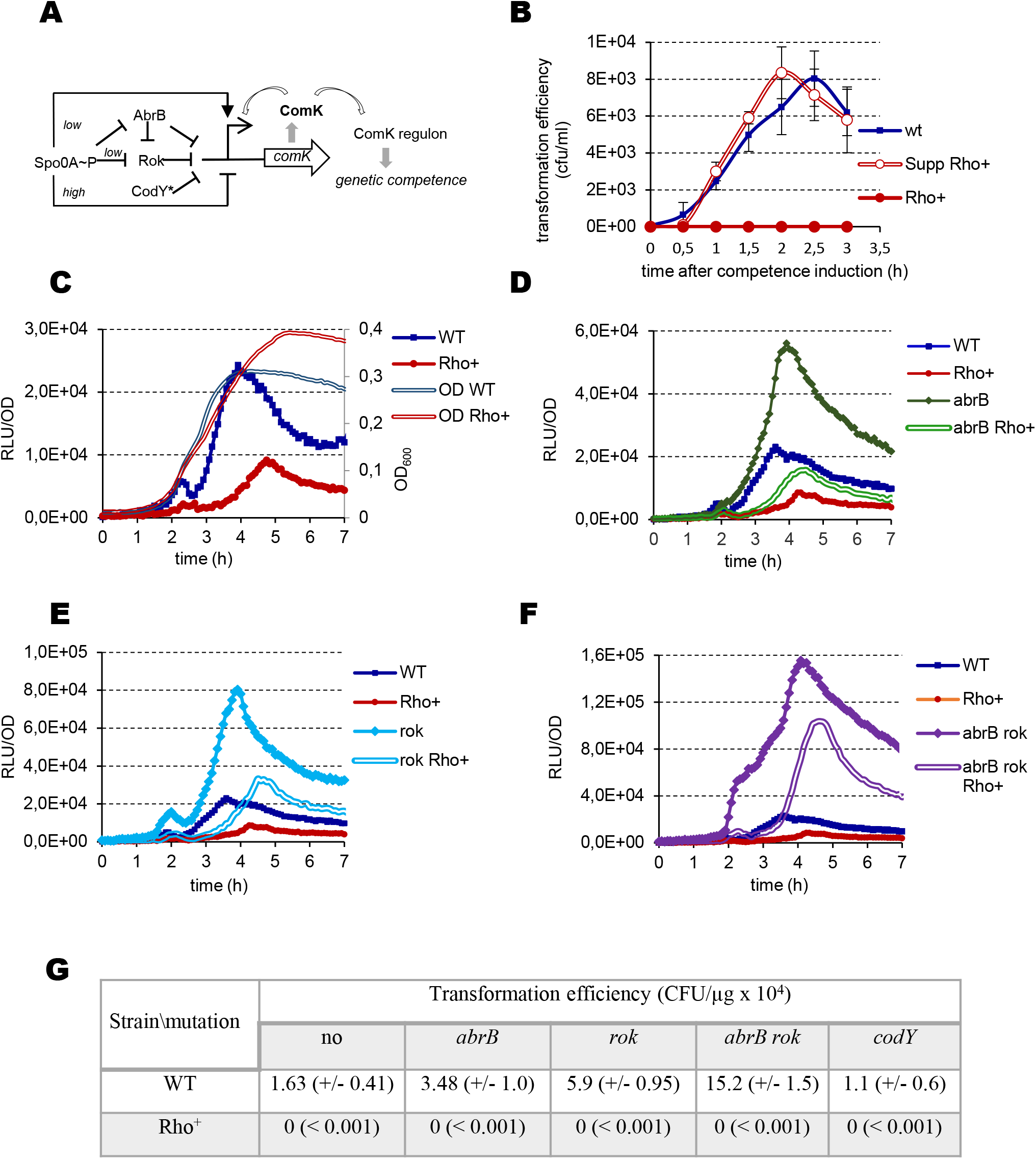
*B. subtilis* Rho^+^ exhibits competence–negative phenotype. **(A)** Schematics of P*_comK_* regulation in *B. subtilis* cells. Arrows and bar-headed lines represent positive and negative effects, respectively. **(B)** Kinetics of competence development in *B. subtilis* WT (blue line), Rho^+^ (red line) and Rho^+^_Q146Stop_ (red double-line) strains. Cells grown in a defined rich medium to stationary phase were transferred to the competence-inducing medium (T0) and tested for transformation by homologous genomic DNA over three hours as described in Materials and Methods. The experiment incorporated three biological replicas of each strain and was reproduced three times. Plotted are the mean values and SD from a representative experiment. **(C)** Kinetics of luciferase expression in *B. subtilis* WT (blue line) and Rho^+^ (red line) cells bearing the P*_comK_-luc* transcription fusion and grown in competence-inducing medium. Plotted are the mean values of luminescence readings corrected for OD from four independent cultures of each strain analyzed simultaneously. The double-lined curves depict characteristic growth kinetics of cells measured by OD_600_. (**D, E and F**) Kinetics of luciferase expression in *B. subtilis* mutant strains: (**D**) *abrB* P*_comK-luc_* (green line) and Rho^+^ *abrB* P*_comK-luc_* (green double-line); (**E**) *rok* P*_comK-luc_* (light blue line) and Rho^+^ *rok* P*_comK-luc_* (light blue double-lines); (**F**) *abrB, rok* P*_comK-luc_* (purple line) and Rho^+^ *abrB, rok* P*_comK-luc_* (purple double-line). The indicated mutant pairs were analyzed in parallel with the control parental strains WT P*_comK-luc_* (blue line) and Rho^+^ P*_comK-luc_* (red line). For each strain, data acquisition and processing were performed as in (C). Each strain was analyzed at least three times. The results of representative experiments are shown. (**G**) Inactivation of the known repressors of *comK* does not rescue competence–negative phenotype of Rho^+^ cells. Effect of Rho over-production on transformation efficiency of *B. subtilis* WT and Rho^+^ strains and their respective derivatives carrying single mutations in the *abrB*, *rok, codY* genes or double mutations in *abrB, rok* genes. Cells were transformed by donor genomic DNA after two hours of growth in competence-inducing medium as described in Materials and Methods. Shown are the mean values with SD (in brackets) from two independent experiments each incorporating three biological replicas of each strain.

As shown in Fig 3B, the efficiency of transformation of WT cells by homologous genomic DNA gradually increased during ∼2.5 hours of growth in competence medium and declined later. In the same conditions, Rho^+^ cells remained transformation-deficient all over the experiment. Similarly, it appeared impossible to transform Rho^+^ cells with plasmid DNA (S2 Fig). The primary role of Rho in the competence–negative phenotype of Rho^+^ cells was further confirmed by the WT-like transformation efficiency of the Rho^+^_Q146Stop_ suppressor mutant selected as restoring sporulation (see above; Fig 3B).

To determine whether the competence-negative phenotype of Rho^+^ cells is caused by low expression of the master regulator of competence ComK [71], we followed the activity of the *comK* promoter (P*_comK_*) during growth in competence medium using a P*_comK_-luc* transcriptional fusion [70]. In accordance with previously published data [70], we observed an increasing expression of the P*_comK_* promoter in WT cells up to the entry into the stationary phase (Fig 3C). In the same time, P*_comK_* activity was reduced about three-fold in Rho^+^ cells compared to WT (Fig 3C), although remained higher than the basal expression level observed in the *spo0A* mutant (S3 Fig). Thus, *comK* expression in Rho^+^ cells appears insufficient to assure a threshold level of ComK required for competence induction [72].

In exponentially growing *B. subtilis* cells, transcription of *comK* is repressed by AbrB, Rok, and CodY [32, 73, 74]; (Fig 4A) and is activated by the raise of Spo0A∼P, which also relieves the AbrB- and Rok-mediated repression thus opening a temporary “competence window” [70, 75]. Considering the low levels of Spo0A expression in Rho^+^ background in DS medium (see above), it was plausible that Spo0A-mediated de-repression of *comK* was inefficient in Rho^+^ cells. Thus, we attempted to increase expression of *comK* by inactivating its known repressors. Introduction of single *abrB* and *rok* mutations in WT cells increased activity of the P*_comK_* promoter about two- and three-fold, respectively (Fig 3D and E), and simultaneous inactivation of both repressors synergistically stimulated *comK* expression (Fig 3F). Concordantly, we observed the increased transformation efficiencies of the mutants in our test for the competence state (Fig 3G). Inactivation of *abrB* or *rok* genes in Rho^+^ cells also led to the de-repression of P*_comK_*, close to or above WT levels, respectively (Fig 3D and E), and the combination of both mutations led again to a synergetic five-fold increase of *comK* expression compared to WT cells (Fig 3F). However, despite the strong stimulation of *comK* expression Rho^+^ cells mutated for *rok* and *abrB* remained non-transformable (Fig 3G).

**Fig 4.**
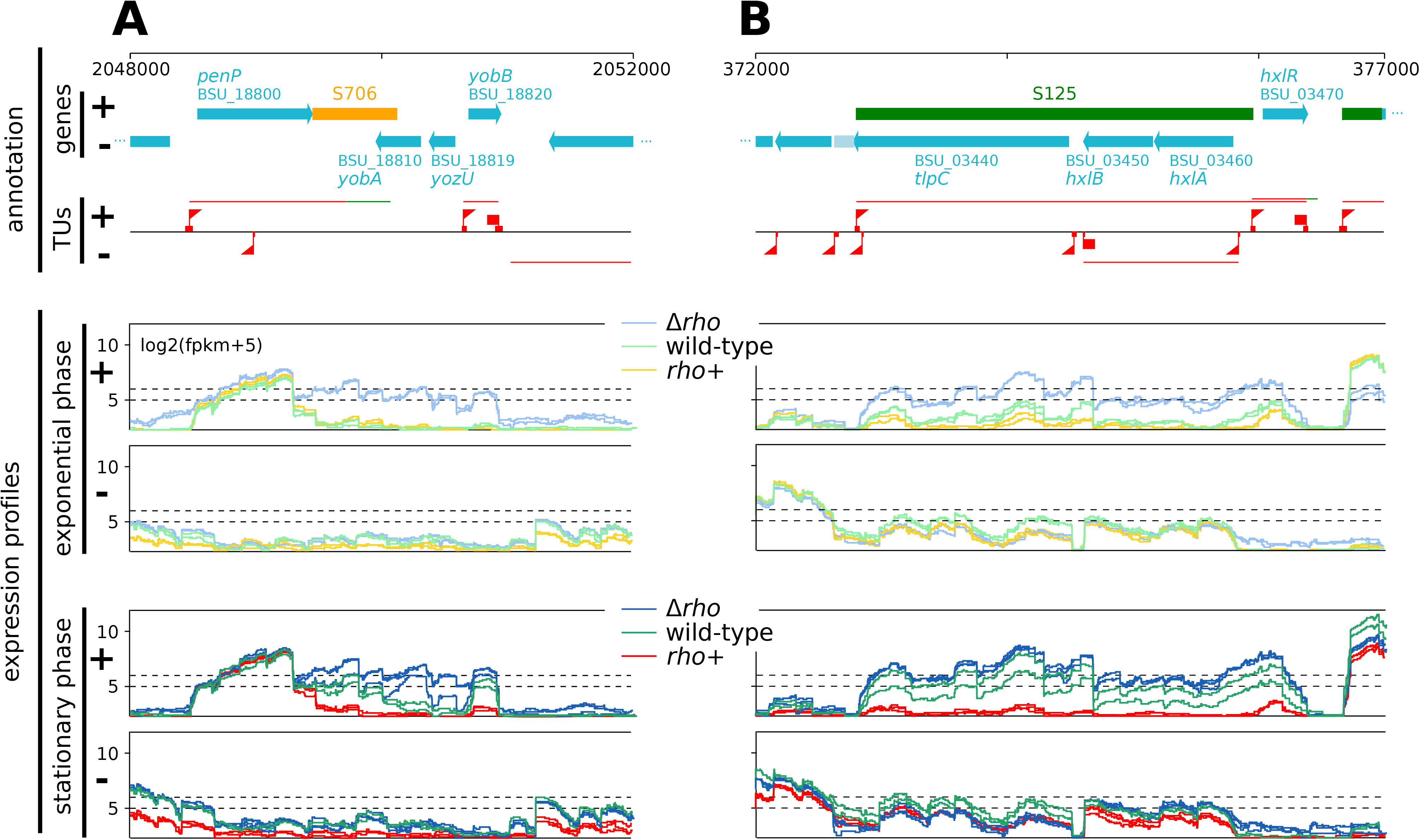
Rho-mediated control of pervasive transcription. Examples of expression profiles of *B. subtilis* WT, Rho^+^ and Δ*rho* strains at two loci (**A** and **B**) as measured by RNAseq in exponential (middle panel) and stationary (bottom panel) growth phases for both strands of the genome (+ and -). The whole genome can be browsed at http://genoscapist.migale.inrae.fr/seb_rho. The top panel presents the structural organization of the region including the GenBank annotation, S-segments and transcription units (TUs) as determined from a compendium of WT expression profiles by Nicolas *et al*. (2012). Triangle and square flags positioned on the TU lane (red) represent identified abrupt transcriptional up- and down-shifts often associated with promoter/terminator activity. The colors of expression profiles (middle and bottom panels) distinguish strains and growth phases: WT (light and dark green lines for exponential and stationary phase, respectively), Rho^+^ (orange and red) and Δ*rho* (light and dark blue). Inactivation of Rho in Δ*rho* mutant leads to the mRNA extension of the 3’UTR of *penP* gene (S706, antisense of *yobA-yozU*, sense of *yobB*) (**A**) and increases the expression of the asRNA (S125, antisense of *tlpC-hxlB-hxlA,* sense of *hxlR*) from its own promoter (**B**). Opposite effects are observed in Rho^+^. While Δ*rho* and Rho^+^ profiles are clearly distinguished in both conditions, the WT is intermediate with a position closer to Rho^+^ in exponential phase and to Δ*rho* in stationary phase (*i.e.* consistent with the decrease of Rho abundance upon transition to the stationary phase in WT).

Repression activity of CodY does not depend on Spo0A∼P as it relies on the nutrient and energy cellular status [30, 32]. To evaluate the significance of CodY-mediated regulation of *comK* expression in the context of high Rho amount, we compared the effect of the *codY* mutation on P*_comK_-luc* activity in WT and Rho^+^ cells. In our experiments, the *codY* mutation reduced the growth rate of both strains in the competence-inducing medium, which explains a delayed induction of P*_comK_* compared to CodY^+^ cells, and had a variable effect on its activity (S4 Fig). We attribute this discrepancy to some uncontrolled fluctuations in the nutrient content between the experiments. The relatively small effect of the *codY* mutation on de-repression of *comK* was observed previously [72]. More importantly, the expression of the *comK* gene in Rho^+^ cells mutated for *codY* always remained below the WT level (S4 Fig). Not surprisingly, Rho^+^ *codY* mutant strain appeared non-transformable (Fig 3G).

Altogether, these results show that a steady high level of Rho caused complex and efficient repression of the competence transcription factor ComK. They also pinpoint the existence of other road-blocks acting downstream of ComK which contribute to the loss of genetic transformation in Rho^+^ cells.

### Comparative transcriptome analysis of *B. subtilis* WT and Rho^+^ strains

To gain deeper insight into the origins of competence- and sporulation-deficient phenotypes of Rho^+^ cells and to reveal other modifications of the transcription conceivably caused by stable expression of *rho*, we performed comparative RNAseq transcriptome analyses of *B. subtilis* WT, Rho^+^ and Δ*rho* strains grown in LB medium. Two time points corresponding to the mid-exponential and early stationary phase were selected for this comparison (Materials and Methods).

On RNAseq data, we conducted differential expression (DE) analyses of sense and antisense strands of the native transcription regions (TRs) composed of 4,292 Genbank-annotated genes and 1,583 other TRs (so called “S-segments”) comprising sense and antisense RNAs (asRNAs) identified in WT from a large collection of expression profiles [6]. In addition to the DE analysis whose results are detailed in S2 Table, we used Genoscapist [76] to set-up a web-site for online interactive exploration of strain- and condition-dependent transcriptional profiles up to single-nucleotide resolution; these profiles are illustrated in Fig 4 and can be accessed at http://genoscapist.migale.inrae.fr/seb_rho/.

The RNAseq data obtained for the Δ*rho* mutant confirmed elevated levels of antisense transcription (Fig 5A, S2 Table) seen in previously published *B. subtilis* Δ*rho* transcriptomes [6, 8]. Reciprocally, we observed a global down-regulation of the antisense transcription in Rho^+^ cells compared to WT (Fig 5A; S2 Table), which is consistent with the well-established role of Rho in the suppression of antisense transcription. Namely, we detected that transcription of the antisense strands of 338 GenBank-annotated genes was down-regulated (log2 Rho^+^/WT ≤ −1; q-value ≤ 0.05) in the exponentially growing Rho^+^ cells, and this trend was even more pronounced in the stationary phase where antisense transcription of 1,550 genes was down-regulated (Fig 4, Fig 5A and S2 Table). In particular, out of the 90 S-segments expressed in WT during the stationary phase (cutoff expression level of log2(fpkm+5)≥5) and previously documented as antisense transcripts [6], 58 S-segments were down-regulated in Rho^+^ (log2 Rho^+^/WT≤ −1; q-value ≤ 0.05). Nevertheless, only a small fraction of the down-regulated antisense transcripts in Rho^+^ cells are expressed in WT at levels that would be considered relevant for classical genes (e.g. 24/338 and 96/1,550 for cutoff expression level of log2(fpkm+5)≥5 in Fig 5A). This observation is consistent with relatively low levels of antisense transcription in WT bacteria [77]. It explains that only a minority of the genes that we report here as with decreased antisense transcription in Rho^+^ were previously documented as being subject to antisense transcription in WT [6].

**Fig 5.**
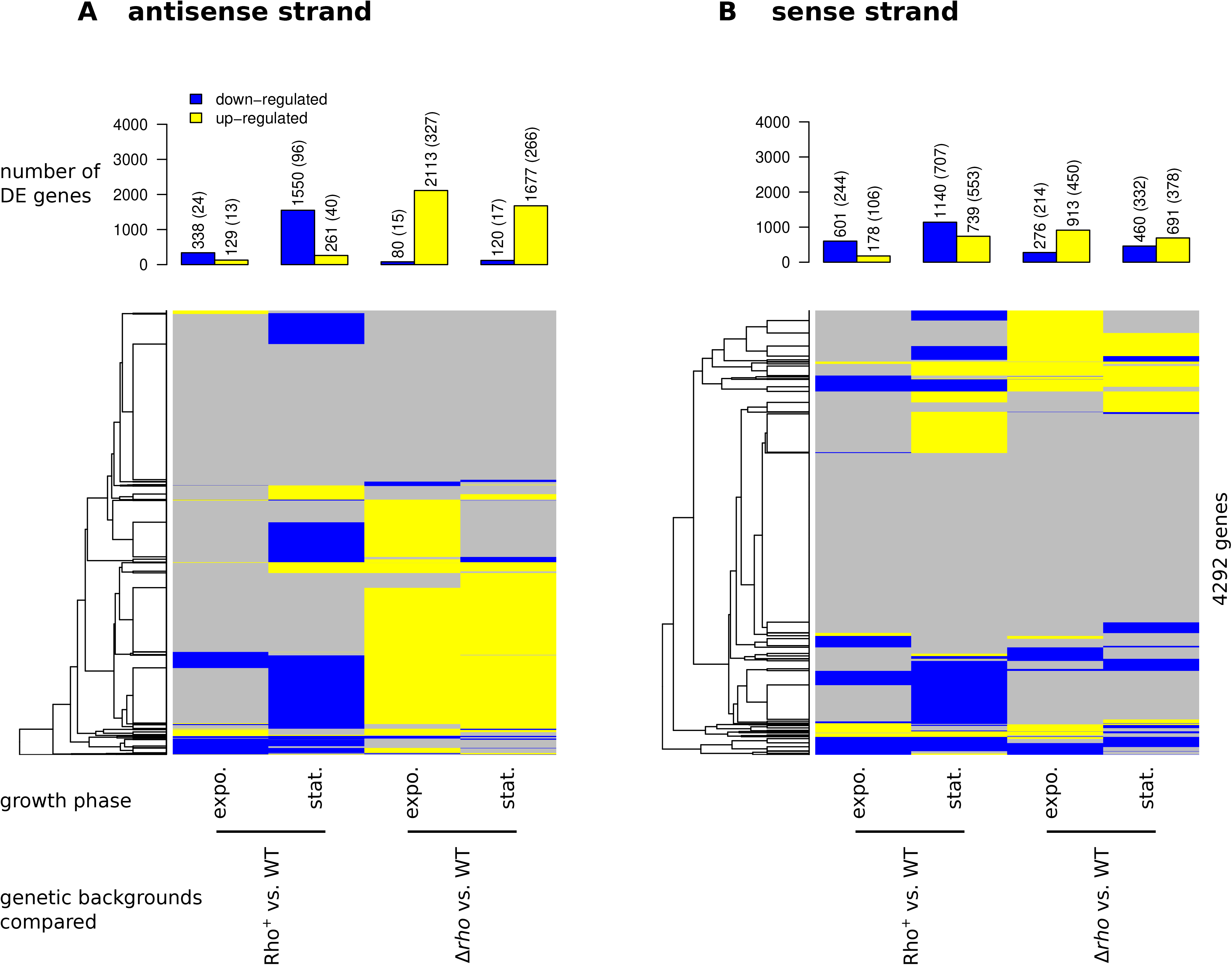
Graphical summary of differential sense and antisense expression (DE) in Rho^+^ and Δ*rho* across strands and growth phases. DE compared to WT (q-value≤0.05 and |log2FC|≥1) is shown for the antisense (**A**) and sense (**B**) strands of the 4,292 AL009126.3-annotated genes. Barplot representation of the numbers of DE genes: numbers reported above each bar correspond to the total of DE genes and, between parentheses, to the subset exhibiting a minimal expression of log2(fpkm+5) ≥5 in one of the two compared genetic backgrounds. Heatmap highlighting overlaps between these sets of DE genes: up-regulated genes in yellow, down-regulated genes in blue, other genes in gray. Left-side of each heatmap: average-link hierarchical clustering tree based on pairwise distance between genes (L1-norm after encoding down-regulation and up-regulation as −1 and 1, 0 otherwise).

Examination of transcriptional profiles along the genome revealed modifications in Rho^+^ cells that are typical for enhanced termination of transcription at a number of weak intrinsic and Rho-specific terminators, preventing read-through transcription often in antisense of downstream genes (Fig 4A). Further supporting this observation of enhanced termination in Rho^+^ cells, we counted that 57 out of 107 S-segments previously described [6] as resulting from a partial termination or exhibiting a 3’ extended mRNA in WT, displayed a decreased expression level in Rho^+^ (log2 Rho^+^/WT ≤ −1).

Over-expression of *rho* also caused considerable modifications of the sense-strand transcription (Fig 5B and S2 Table). However, in contrast to antisense transcription for which inactivation and over-expression of Rho mediated globally opposite effects (up-regulation in Δ*rho* vs. down-regulation in Rho^+^), sense transcription of the Δ*rho* and Rho^+^ strains differed from WT by specific patterns of up- and down-regulations on regions of low and high expression (Fig 5B). This is consistent with changes of the sense-strand transcription caused by a combination of direct effects downstream of Rho-dependent termination sites and indirect effects resulting from their propagation into regulatory cascades, as already described for Δ*rho* [8]. With 739 up-regulated Genbank-annotated genes detected in the comparison Rho^+^ vs. WT in stationary phase, out of which 553 resulting in log2(fpkm+5)≥5; the increased level of Rho in the stationary phase apparently caused the greatest number of indirect effects.

To be able to retrace the propagation of effects into regulatory cascades, we examined the correlation between DE and known regulons and functional categories as defined in *Subti*Wiki database ([78]; http://subtiwiki.uni-goettingen.de). The complete list of statistically significant associations for up- and down-regulated genes in Rho^+^ and Δ*rho* (Fisher exact test p-value ≤ 1e-4) is presented in S2 and S3 Tables, and was further used to investigate alterations of gene expression caused by steady high level of Rho in *B. subtilis* cells. The three strongest statistical associations between regulons and DE gene sets for the comparison Rho^+^ vs. WT in the stationary phase were for AbrB, CodY, and the stringent response regulons (p-value ≤ 1e-12, S3 Table) which are all three known as global regulatory pathways governing the transition to the stationary phase.

Taking into account the pronounced transcriptional changes induced by Rho over-expression in the stationary-phase cells, and the phenotypes of Rho^+^ strain described above, we examine below more closely the expression of genes controlled by ComK, AbrB, and CodY, and the stringent response.

### Suppression of the ComK regulon in Rho^+^ cells

As presented above, over-expression of Rho led to the inhibition of the *comK* promoter activity. In line with this, the amount of the *comK* transcript was significantly reduced in Rho^+^ compared to WT under exponential and stationary conditions (four- and 15-fold, respectively) (S2, S3 Tables and Fig 6A, B). Thus, transcriptome analysis confirmed that the P*_comK_* promoter was not de-repressed upon entering the stationary phase in Rho^+^ cells. Since ComK activates its own transcription, insufficient ComK amount would impede a positive feedback autoregulation of *comK* and activation of the ComK-controlled genes (Smits *et al*., 2005). Indeed, the 34 out of 60 genes belonging to the ComK regulon were down-regulated more than two-fold upon entry into the stationary phase; expression of 20 of them was reduced already during the exponential growth (Fig. 6A, B; S2 and S3 Tables). The strongest transcriptional decrease was detected for the genes involved in binding and uptake of DNA (van Sinderen *et al*., 1995): the *comC*, *comEA-EB-EC*, *comFA-FB-FC*, and *comGA-GB-GC-GD-GE-GF-GG-spoIIIL* operons with a maximum reduction for the *comGC-GD-GE-GF* genes (more than 300-fold; S2 Table). None of the genes involved in DNA recombination and repair (*recA, addAB, sbcD, ssbA, radA* and *yisB*) were repressed in Rho^+^ cells during either exponential growth or stationary phase. While the weak dependence of the *addAB* genes on ComK activity was noticed previously [79–81], the stable expression of other genes might be explained by the presence of additional regulatory elements.

**Fig 6.**
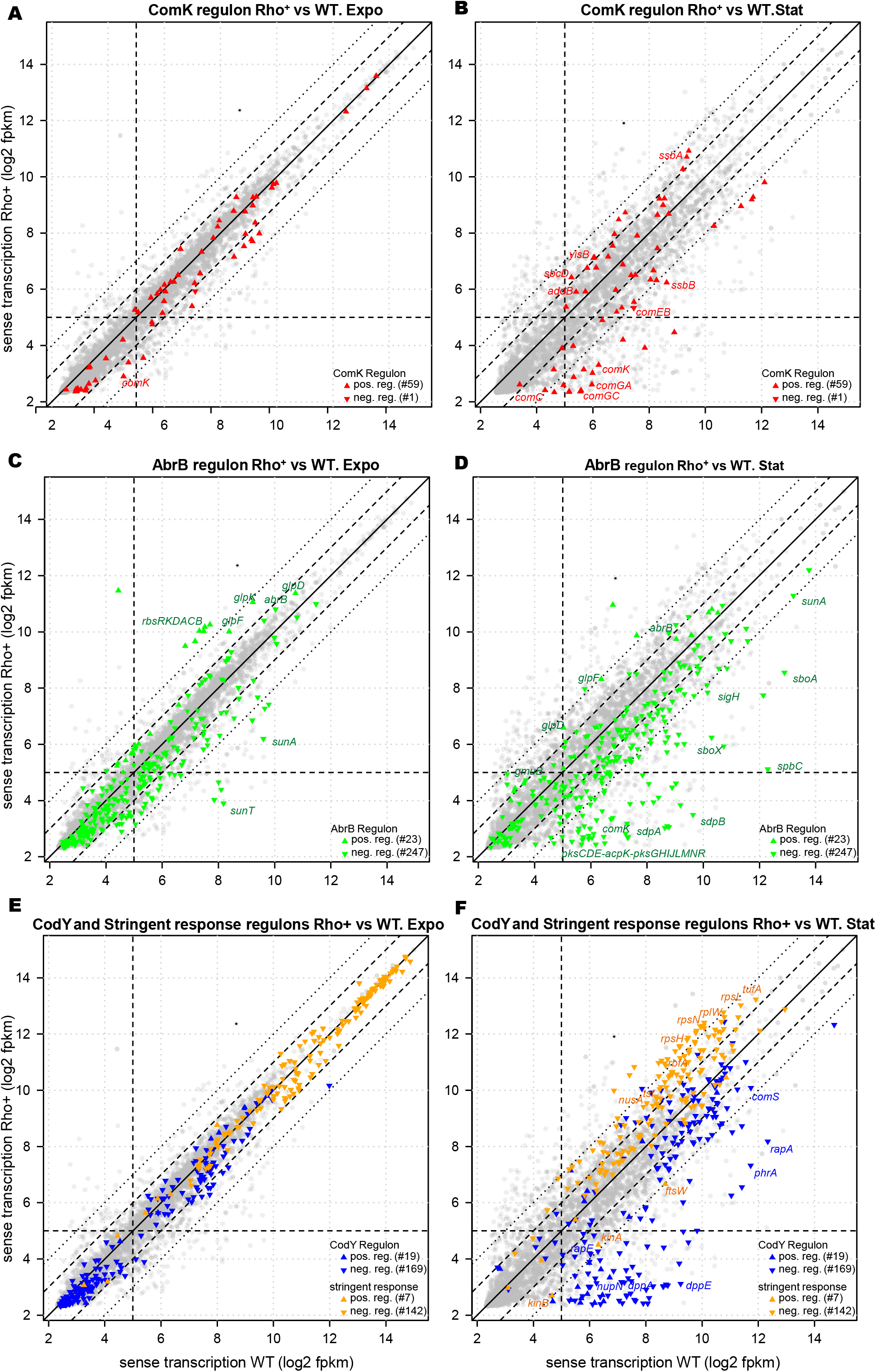
Differential expression of the ComK, AbrB, CodY and stringent response regulons in Rho^+^ strain across exponential growth and stationary phase. Expression changes for the each gene from the ComK regulon (**A** and **B**), AbrB regulon (**C** and **D**), CodY and the stringent response regulon (**E** and **F**) by comparing *B. subtilis* Rho^+^ and WT strains under conditions of exponential growth (A, C, E) and stationary phase (B, D, F). Each point (red for the ComK regulon, green for the AbrB regulon, blue for the CodY regulon, yellow for the stringent response regulon, and gray for genes outside of the mentioned regulons) represents one of the 4,292 AL009126.3-annotated genes. Gene names mentioned in the text are indicated.

In agreement with our previous results [6, 8], no significant differences were found for the expression patterns of the ComK-controlled genes between WT and Δ*rho* strains grown either exponentially or stationary (S6 Fig A, B; and S2, S3 Tables).

### Rho attenuates de-repression of the AbrB-controlled genes

The amount of *B. subtilis* transition state regulator AbrB decreases rapidly upon entering the stationary phase, causing subsequent de-repression of the AbrB regulon of genes [21]. In Rho^+^ cells, the *abrB* gene was up-regulated two-fold in the exponential- and stationary-phase cells (Fig 6C, D and S2, S3 Tables). This most probably resulted from the inefficient activation of Spo0A (see above), which is responsible for the *abrB* repression [25, 26]. In addition, the observed three-fold decrease in *rnaC* sRNA can contribute to stabilization of the *abrB* mRNA in Rho^+^ [27].

Accordingly, 36% and 62% of the negatively controlled genes from the AbrB regulon were significantly down-regulated in Rho^+^ cells compared to WT during exponential growth and in the stationary phase, respectively (Fig 6C, D; and S2, S3 Tables). For example, while the operons encoding antimicrobial compounds (*pksCDE-acpK-pksFGHIJLMNR*, *ppsABCDE*, *sunA*, *sboA-albG* and *bacAF*) (81 Strauch *et al*., 2007) were effectively activated in the stationary WT cells, their expression remained at a lower level in Rho^+^ (Fig 6C, S2 Table). The strong decrease in transcription was also found for the *sdpABC* and the *skfABCEFGH* operons encoding SdpC sporulation delay toxin and the SkfA killing factor, respectively [83]; (Fig 6C, S2 Table). Transcription of the *sigH* gene, which is negatively controlled by AbrB [84], increased upon entry into the stationary phase in WT cells, but was four-fold lower in Rho^+^ (S2 Table). This effect propagated into the SigH regulon, since almost half of the corresponding genes were down-regulated in Rho^+^ cells compared to WT (S2 Table). In accordance with the results showing inefficient de-repression of negatively controlled genes, we noted that several operons activated by AbrB (the *rbsRKDACB*, *glpD*, *glpFK*, and *gmuBACDREFG*) [84], were upregulated in the exponential Rho^+^ cells compared to WT (S2 Table). Upon entering the stationary phase, the expression of all of them (with the exception of the *rbsRKDACB* operon) exceeded significantly the WT level (S2 Table).

This analysis demonstrated that, being at a high level, Rho attenuates de-repression of the AbrB regulon and thereby reduces the expression of numerous transition state genes essential for the adaptation to unfavorable growth conditions.

### Rho restrains de-repression of the CodY regulon upon entry into the stationary phase

The global transcription regulator CodY controls a number of genes essential for the successful transition from the exponential to stationary phase in accordance to the nutrient and energy cellular status [30, 86]. During the exponential growth, all genes repressed by CodY are transcribed at a low level until the cells reach the stationary phase when the activity of CodY starts to decline. In line with this, our analysis showed that over 90% of the CodY-controlled genes were reliably repressed in WT cells during the exponential growth in the rich LB medium compared to the stationary phase (S2 Table). In Rho^+^ cells, this repression was not wholly taken off during the stationary phase for more than 66% of these genes (Fig 6E and 6F; S2 and S3 Tables). Most of them encode proteins involved in amino acid metabolism and are controlled by one or more additional regulators (e.g., AbrB, TnrA or AhrC) responding to other intracellular and/or environmental signals. Others genes, like the *dppA-E* operon encoding a dipeptide permease or the *nupN-Q* operon encoding the guanosine transporter, are under the sole control of CodY. Expression of these genes was strongly decreased in Rho^+^ cells compared to WT (from 25- to 190-fold for *dppA* and *dppE*, respectively, and 36-fold for *nupN-O* genes; S2 Table).

Under nutrient-rich conditions, CodY prevents the development of competence and sporulation [30, 32]. Besides the direct negative effect on the genetic competence through repression of the *comK* promoter, CodY controls negatively the *srfAA-AB-comS-AC-AD* operon, which encodes an essential component of the competence activation pathway, ComS [87]. Transcription of the *comS* gene was significantly down-regulated (more than three-fold, S2 Table) in the stationary Rho^+^ cells compared to WT. CodY controls directly several genes essential for the initiation of sporulation, including the *kinB* gene. In the stationary-phase Rho^+^ cells, expression level of the *kinB* gene was decreased 14-fold (S2 Table). However, we attributed this effect mainly to the improved Rho-dependent intragenic transcription termination of *kinB* [8]. Transcription of the *phrA* and *phrE* genes encoding the regulatory peptides of the Spo0F-specific phosphatases RapA and RapE was repressed ∼27- and nearly three-fold, respectively. All these changes fit with the observed competence- and sporulation-deficient phenotypes of Rho^+^ strain.

We noted that genes strongly down-regulated in Rho^+^ cells fall into the clusters of genes associated with the strongest CodY-binding sites, which were shown to be repressed at an intermediate concentration of active CodY [88, 89]. Thus, we suggested that in the stationary-phase Rho^+^ cells, growth-dependent inactivation of CodY is delayed, and CodY retains the ability to control gene expression. The intermediate levels of active CodY are known to increase the activity of promoters jointly repressed by CodY and another transcriptional factor, ScoC, via feed-forward regulatory loop [90, 91]. Indeed, the *braB* gene and the *oppA-B-C-D-F* operon known to be under the interactive CodY-ScoC regulation were up-regulated in Rho^+^ cells (five- and three-fold, respectively; S2 Table). CodY-controlled gene expression pattern was not significantly modified in the Δ*rho* mutant strain (S2 and S3 Tables, S6 Fig).

Thus, transcription analysis indicates an ineffective inactivation of CodY in Rho^+^ cells entering the stationary phase compared to WT cells. Considering that the modification of CodY activity is caused by changes in intracellular pools of GTP and/or BCAA [30, 34, 92], we suggested that Rho modifies the content of activated CodY through noticeable alterations in the availability of CodY effectors.

### Expression patterns of the stringently controlled genes in WT and Rho^+^ cells are significantly different

Since in *B. subtilis*, CodY activity and the stringent response are tightly linked [36], inefficient de-repression of the CodY regulon in stationary Rho^+^ cells should be associated with altered expression of stringently controlled genes. This prompted us to compare transcription patterns of the stringently regulated genes in WT, Rho^+^ and Δ*rho* strains.

We found no significant difference in the expression of the stringent regulon genes in the exponentially growing WT and Rho^+^ cells (Fig 6E, S2 and S3 Tables). As expected, transcription of the negatively regulated stringent response genes was decreased in WT cells entering the stationary phase. Of these genes, 87% (124 out of 142) showed at least a two-fold decrease in the expression levels compared to the exponential phase (S2 and S3 Tables). In accordance with the former transcriptome analysis of the stringent response [93], genes encoding the components of translational apparatus, including ribosomal proteins (r-proteins) and translation factors were considerably repressed. Of these genes, 91% showed at least a four-fold decrease in their expression levels, while 62% of them were down-regulated more than 10-fold (S2 Table). Conversely, the *kinA*, *kinB*, *ftsW* and *pycA* genes previously shown to be under positive stringent control [94, 95] were up-regulated from four- to eight-fold. These values fit well the changes in the expression of the stringent regulon genes detected earlier in WT cells [6].

The transcription pattern of the stringent regulon was remarkably different in the stationary Rho^+^ cells (Fig 6F, S2 and S3 Tables). In total, 60% of genes negatively regulated by stringent response (86 out of 142) were transcribed from two- to 9-fold more efficiently in Rho^+^ cells than in WT. Almost all genes encoding r-proteins were significantly up-regulated (from two- to five-fold) in Rho^+^ cells compared to WT. The relative increase in transcription of genes encoding the translation factors (*tufA, tsf, fusA, efp, frr, fusA, infA, infB, infC, rbfA*) varied from two- to four-fold. In addition, a similar increase was observed for genes involved in RNA synthesis and degradation (e.g*., nusA*, *nusB*, *rnc*, and *rpoA*). On the contrary, the *kinA*, *kinB* and *ftsW* genes positively regulated by the stringent response [94, 95] were less efficiently transcribed in Rho^+^ cells (Fig 6F and S2 Table). The expression level of the *hpf* gene, which encodes a ribosome hibernation-promoting factor and is considered as a reporter for the activation of stringent response [96], was reduced two-fold in Rho^+^ compared to WT (S2 Table).

We noticed also that some genes whose transcription in response to starvation was shown to be changed in a Rel-independent manner [93] had the opposite behavior in WT and Rho^+^ cells. Indeed, while most of the genes encoding aminoacyl-tRNA synthetases were repressed in the stationary WT cells, their transcription slightly increased in Rho^+^. In accordance with the previous analyses [93, 97], we observed a decrease in transcription of genes involved in purine biosynthesis in the stationary WT cells. In contrast, the *purE-K-B-C-S-Q-L-F-M-N-H-D* operon genes were expressed from three- to six-fold and the *xpt-pbuX* genes up to 20-fold higher in Rho^+^ cells compared to WT (S2 Table).

Comparison of WT and Δ*rho* mutant strains did not reveal any global changes in the transcription of the stringent regulon genes in these growth conditions (S6 Fig and S2, S3 Tables).

Overall, the present analysis revealed significant differences between the expression patterns of the stringent regulon in Rho^+^ and WT cells, which suggests that when steadily expressed, Rho restrains activation of the stringent response upon entry into the stationary phase. Following this hypothesis, we assessed the physiological consequences of *rho* over-expression on characteristic phenotypes achieved through induction of the stringent response in *B. subtilis*.

### Rho^+^ strain exhibits a modified cell morphology and decreased stationary-phase survival

In *B. subtilis* as well as in other bacteria, induction of the (p)ppGpp synthesis under nutrient limitation and activation of the stringent response at stationary phase causes cell size reduction [98–102]. In accordance, the (p)ppGpp-deficient cells are longer than WT cells, which correlates with an altered expression of genes involved in cell shape determination and the biosynthesis of fatty acids and cell wall components [93, 99, 101].

Microscopy analysis of the cellular morphology detected no difference between WT and Rho^+^ strains during exponential growth. However, while WT cells were effectively reduced in size and appeared as short rods with the average length of 1.9 ± 0.6 µm upon entering the stationary phase, Rho^+^ cells remained significantly longer (average cell length 3.2 ± 1.0 µm; Fig 7A, B). Consistently with the observed cell size reduction, 53% of genes involved in cell wall synthesis (the gene functional categories as defined in *Subti*Wiki; [78]) were repressed from two- to ten-fold in WT cells after transition to stationary phase (S2 Table). In contrast, most of these genes displayed a higher expression level in Rho^+^ cells than in WT cells, during stationary phase. For instance, the *murE-mraY-murD-spoVE-murG-B* gene cluster involved in the biosynthesis of peptidoglycan precursors [103]; the *cwlO, lytE* and *ftsX* genes encoding lytic enzymes critical for the cell elongation [104]; and the *mreB-C-D* genes responsible for the cell shape determination [105] were expressed up to four-fold higher in the stationary Rho^+^ cells than in WT (S2 Table).

**Fig 7.**
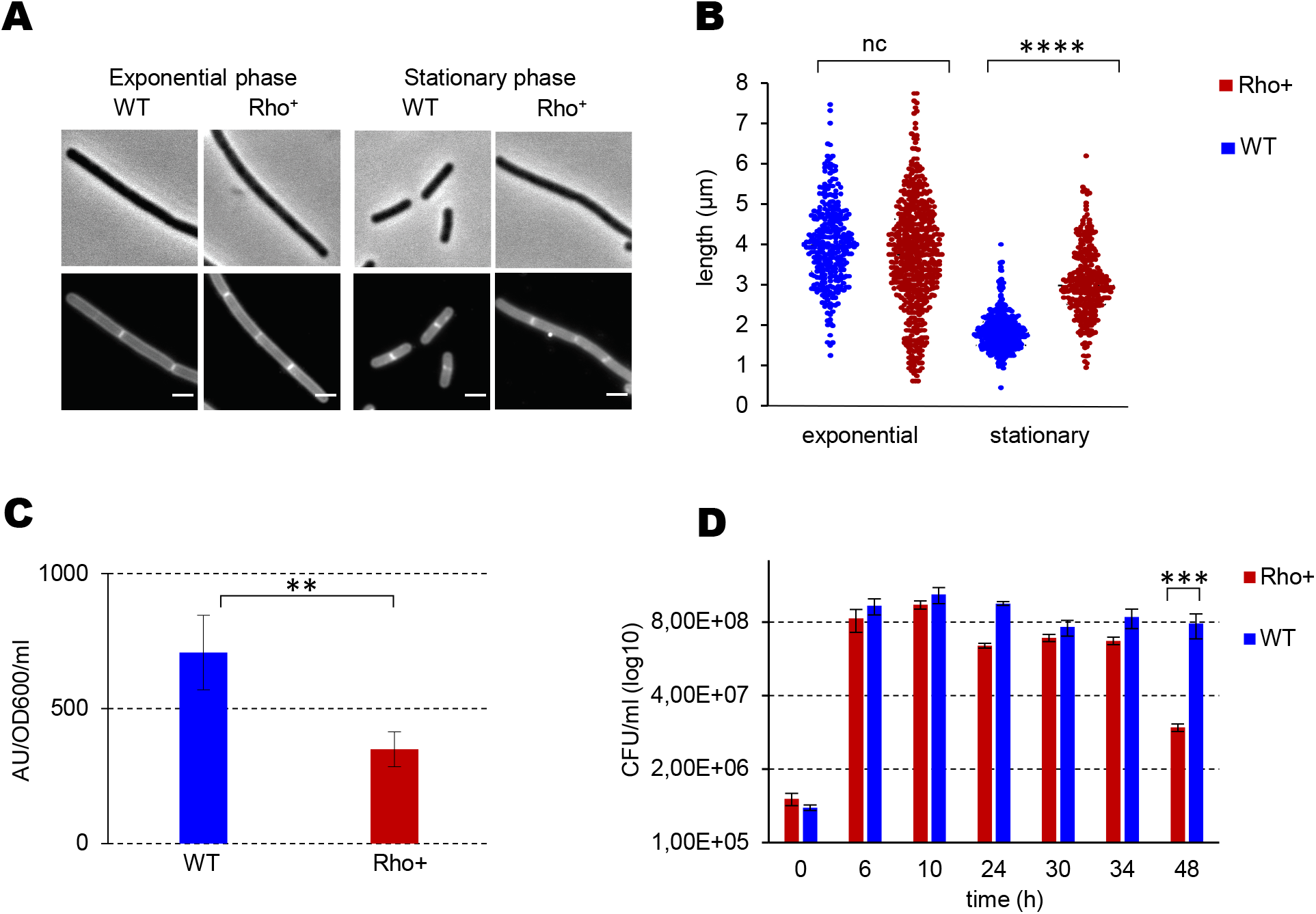
Morphology and long-term survival of *B. subtilis* Rho^+^ cells. Microscopy images (**A**) and cell length measurement (**B**) of *B. subtilis* WT and Rho^+^ under conditions of exponential growth and stationary phase. (A) Phase contrast (upper panel) and fluorescence (lower panel) images of WT and Rho^+^ cells stained with Nile Red at OD_600_ 0.2 and 1.6. Scale bars correspond to 6 µm. (B) The post-acquisition treatment of the images and determination of the mean cell lengths was as described in Materials and Methods. Statistical significance was estimated with a nested t-test, performed with Prism 9 (GraphPad Software, LLC). Note that the plot show the pooled values of the replicates. *P*-values are displayed as follows: **** = P<0.0001; *** = 0.0001<P<0.001; ** = 0.001<P<0.01; * = 0.01<P<0.05; ns = P>0.05; ns, non significant (p>1.0) using a nested t-test. (**C**) Level of ppGpp under stationary phase. *B. subtilis* BsB1 WT and Rho^+^ cells were grown in MS medium supplemented with 0.5% (w/v) of casamino acids to the stationary phase (OD_600_ 1.5). The ppGpp levels were assessed as described in Materials and Methods. Plotted are the mean values and SD from three independent experiments. **, p ≤ 0.01 using a two-tailed t-test. (**D**) Effect of Rho on long-term survival of *B. subtilis. B. subtilis* WT and Rho^+^ cells were grown in LB medium at 37°C with vigorous shaking during 48 hours. At the specified growth time, cells were platted on LB plates, and cell survival in cultures was assessed by the number of viable cells forming colonies (CFU) after 18 hours of incubation at 37°C. Plotted are the average values from three independent experiments each incorporating three biological replicas of each strain. ***, p ≤ 0.001 using a two-tailed t-test.

To assess whether an abnormal size of the stationary-phase Rho^+^ cells correlates with a reduced level of (p)ppGpp alarmon, we compared the ppGpp pools in WT and Rho^+^ cells using high performance liquid chromatography (HPLC). Indeed, we found that Rho^+^ cells grown stationary accumulated about two-fold less ppGpp compared to WT cells (Fig 7C).

Considering the crucial role of the stringent response in the adaptation and survival under starvation conditions [100, 106–109], we next examined whether a steady high Rho amount affects a long-term survival of *B. subtilis*. The growth rate and viability of Rho^+^ cells during the exponential growth in LB medium were identical to those of WT (Fig 1, Fig 7D). However, while almost 50% of WT cells remained viable for at least 48 hours, the viability of the Rho^+^ strain decreased significantly during this time, as estimated by colony formation (Fig 7D). It is noteworthy that a decreased long-term survival of Rho^+^ strain does not depend on its failure to sporulate, since LB medium does not support efficient sporulation, and at 48 hours, lesser than 0.5% of WT cells formed spores.

These results are consistent with RNAseq data, and show that stationary Rho^+^ cells exhibit characteristic features associated with a decrease in intracellular (p)ppGpp levels.

### Rho^+^ strain exhibits phenotypic amino acid auxotrophy

*B. subtilis* mutants deficient in (p)ppGpp production ((p)ppGpp^0^) are characterized by phenotypic auxotrophy for amino acids, in particular, BCAA, threonine, histidine, arginine, tryptophan and methionine, provoked by a deregulation of GTP homeostasis [42, 46, 110]. Therefore, we tested the ability of Rho^+^ cells to form colonies on a minimal MS medium either in the presence or absence of amino acids. Both WT and Rho^+^ strains grew equally well on MS medium supplemented either with casamino acids (CAA) or with eight amino acids listed above (Fig 8A; data shown for MS medium supplemented or not with CAA). However, omission of CAA had a strong inhibitory effect on the colony-forming ability of the Rho^+^ strain, contrary to WT or the Rho^+^ _Q146Stop_ suppressor mutant of sporulation-deficiency (Fig 8A). Therefore, the increased amount of active Rho protein causes phenotypic amino acid auxotrophy of *B. subtilis.* It was previously shown that lowering intracellular level of GTP restores the viability of *B. subtilis* (p)ppGpp^0^ cells in minimal medium without CAA [42, 46, 110]. To analyze whether this was also true for cells over-expressing Rho, we decided to weaken the level of GTP in Rho^+^ strain by mutating *guaB*, the essential gene of the GTP biosynthesis pathway. This was performed by introducing, in the WT and Rho^+^ strains, of the partial loss-of-function point mutations *guaB* T139I and *guaB* S121F, which were previously isolated as spontaneous suppressors of the poly-auxotrophy of *B. subtilis* (p)ppGpp^0^ mutant [42]. Both T139I and S121F *guaB* mutations rescued the auxotrophic phenotype of Rho^+^ strain (Fig 8B), reinforcing potential link between a high Rho content and the shortage of (p)ppGpp.

**Fig 8.**
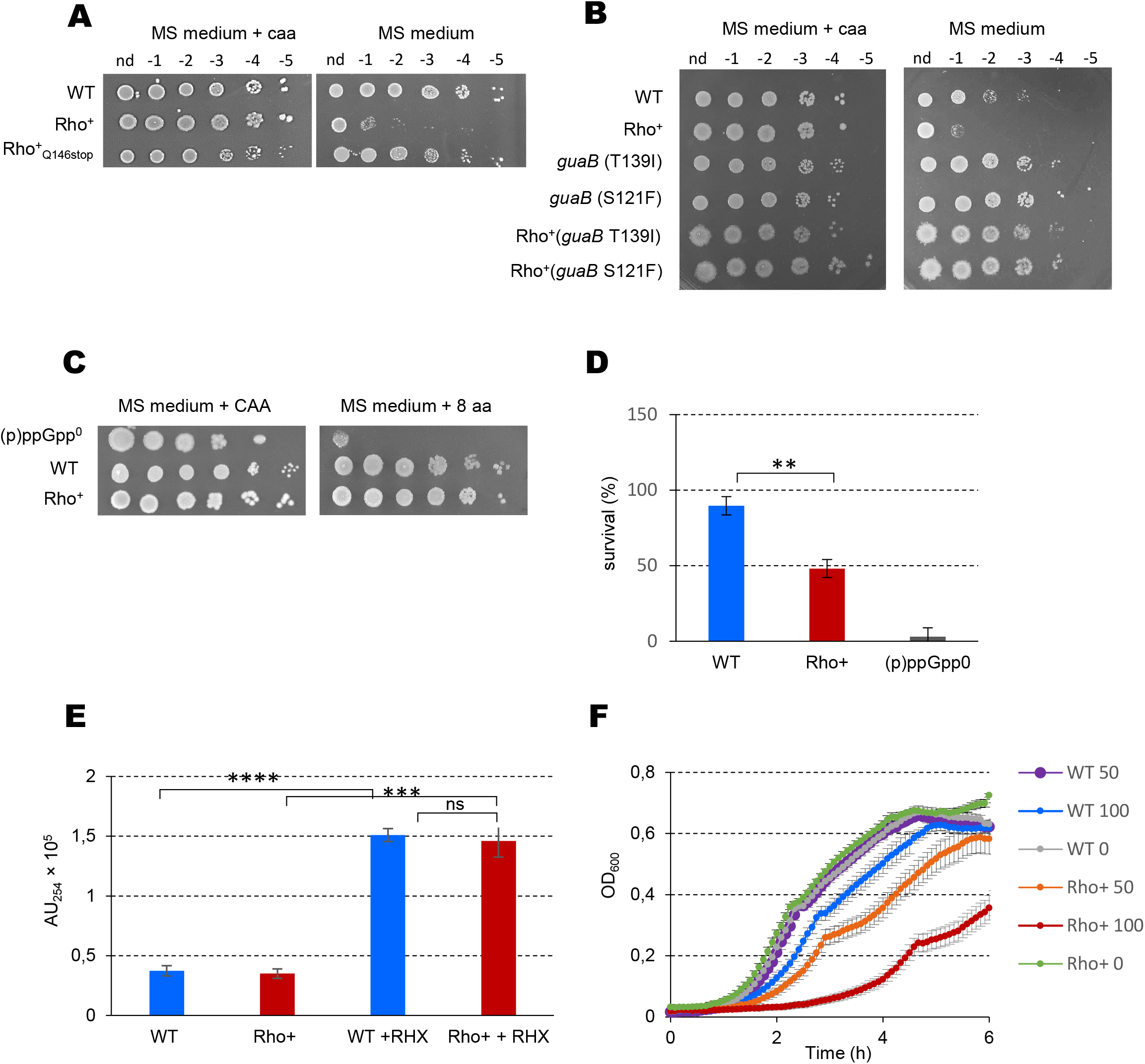
Adaptation of *B. subtilis* Rho^+^ to a sudden nutrient downshift. (**A**) Rho^+^ cells exhibit phenotypic amino acid auxotrophy. *B. subtilis* WT, Rho^+^ and Rho^+^_Q146Stop_ cells growing exponentially (OD_600_ 0.5) in liquid S7 medium containing 0.5% (w/v) of casamino acids (CAA) at 37°C were spotted in serial dilutions on MS agar plates supplemented or not with CAA (0.5%). Plates were incubated at 37°C during 18h before imagining. (**B**) Decreasing of GTP level rescues the auxotrophic phenotype of Rho^+^ cells. *B. subtilis* WT, Rho^+^ and their respective derivative strains carrying *guaB*S121F and *guaB*T139I mutations were cultivated in liquid S7 medium containing 0.5% (w/v) of CAA and tested for the ability to grow in the absence of CAA as in (A). (**C**) Unlike (p)ppGpp^0^ cells, Rho^+^ strain resists to mild nutrient limitations. Isogenic WT, (p)ppGpp^0^ and Rho^+^ strains were grown in liquid MS medium containing 0.5% (w/v) of CAA and spotted in serial dilutions on MS agar plates supplemented either with 0.5% (w/v) of CAA or with 0.05mg/ml of each of the eight amino acids (valine, isoleucine, leucine, threonine, histidine, arginine, tryptophan and methionine). Plates were incubated at 37°C during 18h before imagining. In (A, B and C), the experiments were reproduced at least three times and the representative results are shown. (**D**, **E** and **F**) Rho^+^ cells exhibit altered resistance to arginine hydroxamate (RHX). (**D**) Isogenic *B. subtilis* WT, (p)ppGpp^0^ and Rho^+^ strains were grown to the middle exponential phase (OD_600_ 0.5), treated with 500 µg/ml of RHX for 40 min and plated on LB agar plates. Plates were incubated for 18 h at 37°C before counting viable cells that formed colonies. Strain survival upon sudden amino acid starvation induced by RHX was estimated as the percentage of viable cells after and before the treatment. Plotted are the average values and SD from three independent experiments incorporating three biological replicas of each strain. **, P ≤ 0.005 using a two-tailed t-test. (**E**) Increase of ppGpp level following sudden amino acid starvation. *B. subtilis* WT and Rho^+^ cells were grown in MS medium supplemented with 0.5% (w/v) of CAA to the middle exponential phase (OD_600_ 0.5) and treated or not with 500 µg/ml of RHX. Cells were harvested 20 min after addition of RHX, and ppGpp levels were assessed as described (Materials and Methods). Plotted are the average values and SD from three independent experiments. ****, p ≤ 0.0001; ***, p ≤ 0.001; **, p ≤ 0.01; ns, non-significant (p>1.0) using two-tailed t-test. (**F**) Growth defect of Rho^+^ strain in the presence of RHX. *B. subtilis* WT and Rho^+^ strains were cultivated in LB medium without or with RHX added at concentrations 50 and 100µg/ml in a 96-well microplate. Growth of the cultures was monitored by OD_600_ measurement at the five-minute intervals using a microplate reader. Plotted curves are the average OD reads of two independent cultures of each strain grown in triplicates at each condition. The analysis was performed three times; the results of a representative experiment are presented.

However, Rho^+^ cells did not exhibit all known phenotypes characteristic to the absence of (p)ppGpp. For example, it has been shown that (p)ppGpp^0^ cells adapt poorly to a sudden nutrient downshift, which results in a failure to survive the transition from amino acid-replete medium to amino acid-limited medium [46]. Contrary to (p)ppGpp^0^ cells, Rho^+^ cells propagated in liquid MS medium containing CAA formed colonies at a solid MS medium supplemented with only eight amino acids similarly to WT strain (Fig 8C).

Then we tested whether Rho^+^ cells could survive treatment with a nonfunctional amino acid analog arginine hydroxamate (RHX), an inhibitor of arginyl-tRNA synthesis and a powerful activator of the stringent response. Rapid death upon RHX treatment is a characteristic feature of (p)ppGpp^0^ cells [42, 110]. However, using the same protocol of pulse treatment of cells with RHX [42], we observed a rather minor effect of RHX on the viability of Rho^+^ cells. While (p)ppGpp^0^ cells survive very poorly to sudden starvation provoked by the RHX treatment, the survival rate of the Rho^+^ strain was about 50% compared to 98% of WT (Fig 8D). In addition, HPLC analysis revealed very similar increase of ppGpp pool in the exponentially growing WT and Rho^+^ cells in response to RHX treatment (Fig 8E). Nevertheless, Rho^+^ cells appeared highly sensitive to constant exposure to RHX during growth in rich LB medium or in MS medium supplemented with CAA (Fig 8F).

Taken together, these findings converge to the conclusion that Rho^+^ strain differs markedly from WT by its reduced capacity to synthesize (p)ppGpp and to induce the stringent response under some stressful conditions.

### Rho weakens survival of *B. subtilis* cells under fatty acid starvation and heat stress

It has been shown that (p)ppGpp deficiency caused specifically by inhibition of the synthetic activity of the bifunctional synthetase-hydrolase Rel, results in a high sensitivity to fatty acid starvation and heat stress [101, 111]. Therefore, we analyzed the Rho^+^ strain for these phenotypes.

We assessed the ability of Rho^+^ cells to adapt to fatty acid starvation using cerulenin, an inhibitor of the fatty acid synthesis enzyme FabF [112]. As reported previously [101], treatment with cerulenin did not affect viability of WT cells but appeared highly toxic for the (p)ppGpp^0^ strain. Addition of the drug had a less pronounced but significant inhibitory effect on Rho^+^ cells resulting in an efficient growth arrest and concomitant loss of viability as assessed by colony formation (Fig 9A and B).

**Fig 9.**
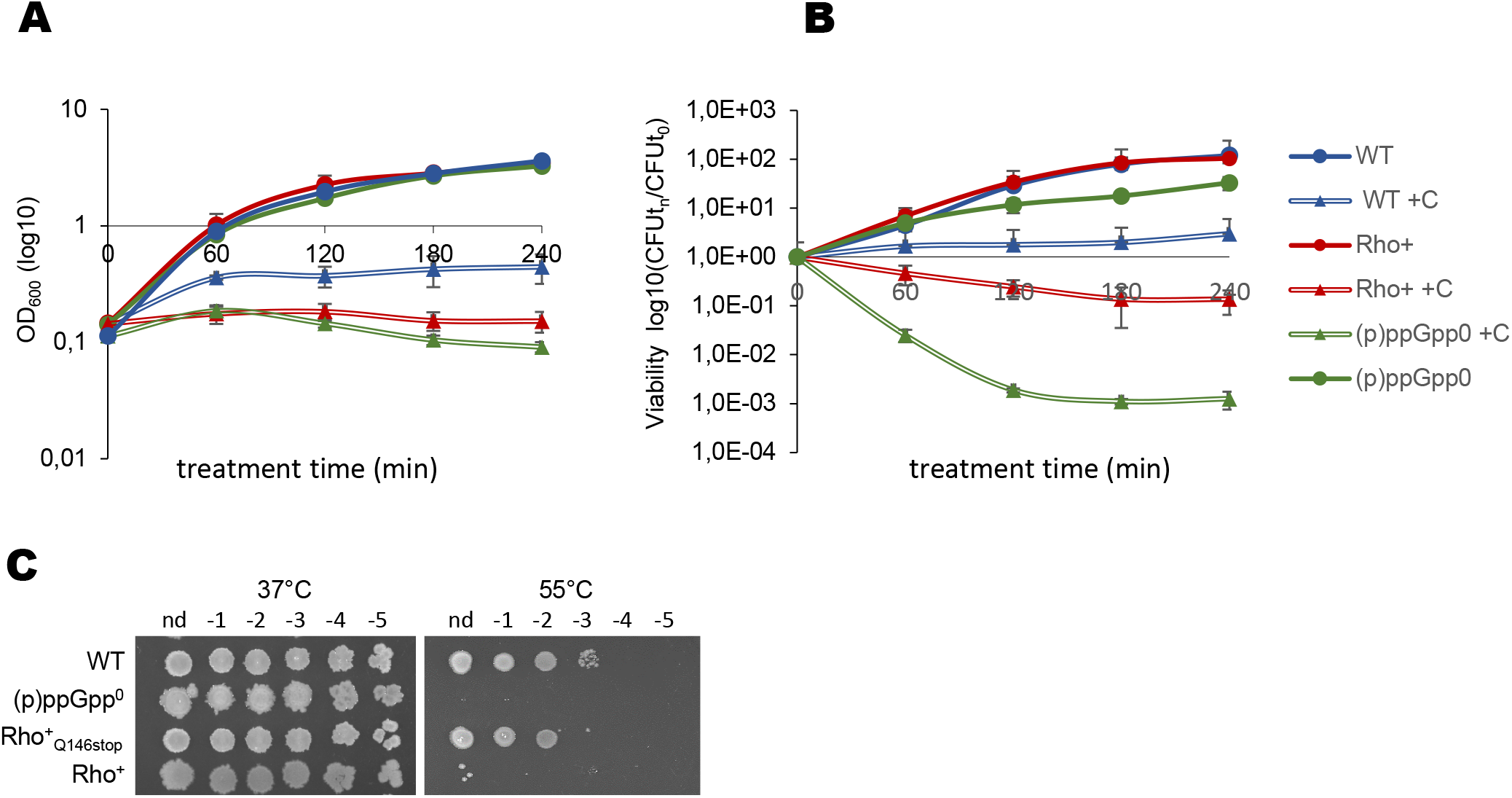
Sensitivity of *B. subtilis* Rho^+^ to fatty acid starvation and heat stress. Growth curves (**A**) and viability (**B**) of *B. subtilis* WT (blue lines), its isogenic (p)ppGpp^0^ (green lines) and Rho^+^ (red lines) strains non-treated (solid lines; circles) or starved for fatty acids by treatment with cerulenin (+ C; double lines; triangles). Cells were grown in LB at 37°C to OD_600_ 0.1, and cerulenin was added to the halves of cultures to final concentration 10µg/ml. Cell survival was estimated by plating bacterial cultures at the indicated time on LB agar and counting of CFUs after 18 h of incubation at 37°C. Each data point in B is the mean of at least three counts. (**C**) Colony formation of *B. subtilis* WT and its isogenic (p)ppGpp^0^, Rho^+^ and Rho^+^_Q146Stop_ (Sup 1) strains at 37°C and 55°C. Cell cultures growing exponentially in LB medium at 37°C were spotted in serial dilutions (from 0 to 10^-5^) on LB agar plates and incubated at 37°C or at 55°C for 18 h before imagining.

Next, we examined the growth capacity of WT and Rho^+^ cells at 55°C, the temperature shown to be non-permissive for both the (p)ppGpp^0^ and synthetase-deficient *relA* mutants [111]. While growth of WT cells platted at solid LB medium was not affected at 55°C, Rho^+^ strain did not form colonies at this temperature (Fig 9C). Furthermore, thermo-sensitive phenotype of Rho^+^ strain allowed us to isolate the mutants able to grow at 55°C. The subsequent analysis of several thermo-resistant Rho^+^ clones revealed mutations of the ectopic *rho* expression unit, similarly to the suppressors of the Rho^+^ sporulation deficiency (S1 Table). These findings confirm that a high level of Rho underlies the heat sensitivity of Rho^+^ cells. Notably, neither *guaB* T139I nor *guaB* S121F mutation did not improve the resistance of Rho^+^ strain to high temperature (S7 Fig), indicating that lowering the cellular GTP level is not sufficient to confer thermo-resistance to *B. subtilis* (p)ppGpp-deficient cells as was shown previously [111].

Taken together, these results demonstrate the involvement of Rho in the control of specific stress survival and suggest that synthetic activity of the main (p)ppGpp synthetase-hydrolase Rel is altered in Rho^+^ cells.

## Discussion

Soil-dwelling bacterium *B. subtilis* adapts to adverse environmental conditions by various survival strategies from the adjustment of metabolic processes via the stringent response to sporulation as an ultimate survival option. Here we provide evidence that the transcription termination factor Rho is involved in the control of the gene regulatory networks that govern adaptation of *B. subtilis* to the stationary phase and thereby in survival under suboptimal environmental conditions.

Previously, using a Δ*rho* mutant we have shown that Rho negatively affects the activity of Spo0A, the master regulator of *B. subtilis* differentiation, by reducing the expression of the major sensor kinase, KinB. Consistently, in the Δ*rho* cells, Spo0A∼P rapidly reaches a high, sporulation-triggering, threshold due to an increased activity of the phosphorelay, which causes accelerated sporulation above the wild type level [8]. We have proposed that Rho-mediated control of the phosphorelay is an essential element of the gene regulatory network centered on Spo0A. In order to maintain the proper rate of sporulation, this control should be released by a programmed decrease of Rho abundance on the onset of sporulation [8].

To assess further the potential regulatory role of Rho, here we used the opposite experimental approach by artificially maintaining *rho* transcription at a relatively high stable level to compensate for the drop of *rho* expression in WT strain upon entering the stationary phase (Fig 1). We show that maintaining stable *rho* expression causes transcriptional reprogramming of *B. subtilis* stationary phase and leads to considerable physiological changes, thereby affecting adaptation of cells to nutrient limitations and cell-fate decision-making to such an extent that it blocks competence development and sporulation.

While inhibition of sporulation due to the repression of *spo0A* transcription at a high Rho level was expected, to some extent, as oppositely mirroring its acceleration in the *rho* mutant cells, loss of genetic competence revealed a novel aspect of Rho-mediated regulation. The master regulator Spo0A∼P plays an essential role in the development of genetic competence by relieving the *comK* gene transcription from the repression by AbrB and Rok [24, 70]. Thus, strong Rho-mediated repression of the *spo0A* gene is consistent with low expression of *comK* and, consequently the ComK regulon.

However, in spite of de-repression of the *comK* promoter in the Rho^+^ *rok*, *abrB* cells, which should provide a threshold ComK level sufficient for activation of the competence genes, this strain retained a competence-negative phenotype manifested by the absence of genetic transformation (Fig 3G). While emphasizing a low Spo0A∼P as a main cause of inefficient expression of the *comK* gene in Rho^+^ cells, these results strongly suggest that a stable high level of Rho also impedes the expression of other gene(s) directly or indirectly involved in the development of genetic competence. There is nothing unusual about this, since many proteins that have been shown to be essential for genetic transformation act independently and downstream of ComK [80, 114–116]. Interestingly, many of these proteins play essential roles in the RNA metabolism [114–116]. As in the case of Rho, their respective roles in the development of genetic competence remain elusive [114, 116]. Further research is needed to identify the potential Rho targets that may be crucial for genetic transformation.

In *B. subtilis,* the role of Spo0A∼P goes beyond the regulation of genetic competence and sporulation. Through the control of the transition state regulator AbrB, Spo0A∼P mediates the de-repression of genes important for adaptation to the stationary phase [21, 85]. Consequently, extended repression of the AbrB regulon observed in Rho^+^ cells (Fig 6D, S2 and S3 Tables) is consistent with a low Spo0A∼P, which appears insufficient for negative regulation of AbrB. Notwithstanding the important role of Spo0A in the cellular adaptive response to nutrient limitations, our results demonstrate that a steady low level of Spo0A∼P is not a sole cause of the complex and coordinated reprogramming of the stationary-phase gene expression in Rho^+^ cells.

Expression of CodY, the second major regulator of the transition to the stationary phase, is independent from Spo0A∼P. The activity of CodY is controlled by the GTP levels, which decrease upon transition to stationary phase due to an increasing synthesis of (p)ppGpp [41–43, 51]. Therefore, the GTP/ (p)ppGpp switch manifests itself in the synchronized activation of the CodY regulon and repression of genes from the stringent regulon, which are required for growth and division [36]. The RNAseq analysis unveils that, in deep contrast with WT strain, in which the amount of Rho decreases upon entering the stationary phase, maintaining Rho at a roughly constant level in Rho^+^ strain impedes de-repression of the CodY regulon and detains the stringent response-related transcriptional changes (Fig 6F, S2 and S3 Tables).

There are no significant differences in the expression levels of genes from the GTP biosynthesis pathways between Rho^+^ and WT strains (S2 Table). This indicates a less efficient accumulation of (p)ppGpp in Rho^+^ cells, rather than an increase in GTP biosynthesis. Concordantly, we detected ppGpp at about twice-lower level in stationary-phase Rho^+^ cells compared to WT and demonstrated that Rho^+^ strain exhibits phenotypic poly-auxotrophy, a hallmark of *B. subtilis* (p)ppGpp-deficiency ([46]; Fig 7C and 8A, B). This growth defect of Rho^+^ strain was rescued by mutations that reduce the synthesis of GTP (Fig 8B), which reflects the restoration of the GTP/ (p)ppGpp balance necessary for the activation of amino acids biosynthetic pathways [46, 110].

In *B. subtilis* cells, accumulation of (p)ppGpp is determined by joint activities of the alarmone synthetases and hydrolases [96, 117–119]. Whereas the expression of both small (p)ppGpp synthetases RelP and RelQ is mainly controlled at the transcriptional level and depends on the growth phase, the bifunctional synthetase-hydrolase enzyme Rel is regulated at the allosteric level [39, 40]. In addition, the activation of RelQ, which is mainly present in a “passive” state, requires (p)ppGpp provided by the bi-functional Rel enzyme [120, 122]. According to the RNAseq analysis, the transcription levels of the *relA*, *relP* and *relQ* genes, and (p)ppGpp hydrolase gene *nahA* (*yvcI*) were similar in Rho^+^ and WT cells (S2 Table). Thus, a decreased level of (p)ppGpp in stationary Rho^+^ cells cannot be caused by changes in the expression of enzymes synthesizing or hydrolyzing (p)ppGpp. In this context, it is important to note that Rho^+^ cells exhibit a thermo-sensitive phenotype, a decreased viability during fatty acid starvation and a reduced long-term survival (Fig 7D and Fig 9). Considering that the bi-functional synthetase-hydrolase Rel is the main source of (p)ppGpp necessary for the survival of *B. subtilis* under these stressful conditions [109, 101, 111], we assume that partially relaxed phenotype of Rho^+^ strain is determined by insufficient accumulation of (p)ppGpp mediated by Rel.

The bifunctional Rel protein can be present in a cell in two alternative states: synthetase-ON/hydrolase-OFF and synthetase-OFF/hydrolase-ON for alarmone synthesis and hydrolysis, respectively [119]. Accordingly, the balance between the synthetase and hydrolase activities of Rel determines the intracellular levels of (p)ppGpp. The Rel-specific synthesis of (p)ppGpp is triggered by ribosomal complexes harboring uncharged tRNA in the ribosomal A-site upon amino-acid starvation [39, 40].

It is known that the enzymatic activity of divers (p)ppGpp synthetic and/or hydrolytic enzymes is modulated by direct interaction with other proteins. In *E. coli*, synthetic activity of bifunctional synthetase-hydrolase SpoT is triggered by the YtfK protein [123] and the acyl carrier protein [124], while its hydrolase activity is stimulated by the anti-sigma factor Rsd [125]. The activity of *E. coli* monofunctional (p)ppGpp synthetase RelA is inhibited by specific interaction with NirD, a small subunit of the nitrite reductase [126]. In *B. subtilis* and Gram-positive pathogen *Listeria monocytogenes*, cyclic-di-AMP-binding proteins DarB and CbpB, respectively, stimulate the synthesis of (p)ppGpp by Rel through direct protein-protein interaction under specific conditions of low intracellular cyclic-di-AMP (c-di-AMP) level [127, 128]. Considering that no difference was found between Rho^+^ and WT cells in the expression of the *ktrA* gene, which is controlled by a c-di-AMP dependent riboswitch ([129]; S2 Table), we concluded that the level of c-di-AMP remains unchanged in both strains under the experimental conditions used. Thus, any particular involvement of DarB into Rel activity in Rho^+^ cells seems unlikely. The late competence protein ComGA interacts with Rel, inhibiting its hydrolase activity, which leads to a temporary increase of the (p)ppGpp pool in competent cells [113]. One can assume that the absence of ComGA due to the inhibition of *comK* expression may contribute to the stabilization of the synthetase-OFF/hydrolase-ON state of Rel enzyme in Rho^+^ cells. However, the previous transcriptional analysis of the *comK* mutant [80] did not reveal changes in gene expression indicative of an altered Rel activity upon entering the stationary phase. In addition, unlike Rho^+^ cells, the *comK* and *comGA* mutants are heat-resistant (S8 Fig), which indicates a sufficient level of (p)ppGpp synthesis mediated by Rel in these strains. Altogether, these data imply that repression of *comGA* cannot underlay a decreased accumulation of (p)ppGpp in Rho^+^ cells.

As a central element of adaptation to various stressful conditions, the stringent response alarmone (p)ppGpp has a strong influence on the cell fate decision-making, regulating the corresponding gene networks at different levels [41, 51, 109, 113, 120, 121, 130].

The (p)ppGpp is involved in the control of genetic competence through the modulation of CodY activity by lowering GTP level [32, 33] and the inhibition of cell growth caused by the ComGA-mediated increase of (p)ppGpp pool [113]. Thus, insufficient synthesis of (p)ppGpp in Rho^+^ cells probably contributes to repression of the *comK* gene by CodY, although a low Spo0A∼P level appears more important for this phenomenon.

A sharp drop in the GTP level is one of the well-known sporulation triggers [41, 51, 130, 131]. It has been shown that *relA* and (p)ppGpp^0^ mutants delay *spo0A* transcription due to a weak activity of the SigH-dependent promoter Ps [52, 132]. In addition, the balance between (p)ppGpp and GTP has a strong effect on transcription of the *kinA* and *kinB* genes, which are under positive stringent control depending on adenine as the transcription initiation nucleotide [94, 95]. Thus, attenuation of (p)ppGpp synthesis provides additional negative regulation of the phosphorelay, contributing to a low level of Spo0A∼P in Rho^+^ cells. That is probably why the artificially increased levels of KinA or KinC kinases, able to trigger sporulation even in nutrient-rich conditions [14, 133], did not completely rescue the sporulation-negative phenotype of Rho^+^ cells (Fig 2E, F).

From this point, it is important to note that the activity of Rel appears crucial throughout the entire pathway of sporulation. The direct evidence for this was provided by the study of Relacin, a potent inhibitor of the Rel-mediated (p)ppGpp production [109]. Authors demonstrated that Relacin strongly inhibited formation of spores regardless the time at which it was added to cells committed to sporulation. In addition, given an important and ever growing number of genes involved in the process of spore formation downstream of the Spo0A phosphorelay [115, 134, 135], we cannot exclude that Rho might negatively regulate some of them.

Taken together, these results show that the transcription termination factor Rho imposes a new layer of control over the stationary phase and post-exponential adaptive strategies. This pinpoints that a *programmed* decrease of Rho levels during the transition to stationary phase is crucial for the adaptation of *B. subtilis* to nutrient starvation: from the adjustment of cellular metabolism and to the activation of survival programs.

Previous studies have shown that in *B. subtilis* and other bacteria, *rho* is negatively auto-regulated through transcription attenuation mechanism at its leader mRNA [5, 136]. In *Salmonella*, the small noncoding RNA SraL was shown to base pair with *rho* mRNA upregulating its expression in several growth conditions [137]. This highlights the importance for cells to regulate the levels of Rho, although the exact mechanism of this control in *B. subtilis* remains to be established.

We propose that in *B. subtilis*, in addition to controlling the Spo0A phosphorelay, Rho participates in the regulation of stationary phase-associated phenomena by tuning the enzymatic ON/OFF balance between the synthetase and hydrolase activities of the bi-functional Rel protein, thereby limiting the (p)ppGpp accumulation upon different stresses.

As a key player of the physiological regulation, (p)ppGpp has been shown to be important for bacterial virulence, survival during host invasion, antibiotic resistance and persistence in both Gram-negative and Gram-positive bacteria [138–141]. Consequently, (p)ppGpp metabolism is currently recognized as a potential target for improving antimicrobial therapy [109, 139, 141]. Albeit the precise molecular mechanism by which Rho delays the accumulation of (p)ppGpp and weakens the stringent response awaits further investigation, the unexpected involvement of Rho in the metabolism of (p)ppGpp should be of particular interest, given the importance of this second messenger for bacterial physiology.

There is increasing evidence for the important role of Rho in controlling various processes connected with (p)ppGpp metabolism (cell fate decisions, virulence and antibiotic susceptibility; [8, 12, 13, 16, 17, 142–144, 147]). Remarkably, mutations of the *rho* gene altering Rho activity were shown to increase adaptation of *E. coli* and *B. subtilis* cells to various stresses and survival under restrictive conditions [145–149].

Taken together, these and the present study highlight the importance of Rho-mediated regulation of genes expression for adaptation to nutrient deprivation and/or other stresses, as well as for the activation of the alternative survival strategies. They unveil Rho as a novel stationary phase regulator and encourage future research.

## Materials and methods

### Bacterial strains and growth conditions

*B. subtilis* strains used in the work are listed in S4 Table. Cells were routinely grown in Luria-Bertani liquid or solidified (1.5% agar; Difco) medium at 37°C. Where indicated, S7 defined synthetic medium [150] containing 50 mM 3-(*N*-morpholino)propanesulfonic acid (MOPS) and supplemented with 0.1% (wt/vol) glutamate, 0.5% (wt/vol) glucose, 0.5% (wt/vol) Casamino Acids (Bacto Casamino Acids), and 0.01% (wt/vol) tryptophan was used. To perform fluorescence microscopy, *B. subtilis* WT and Rho^+^ cells were grown in LB medium to exponential and early stationary phase (optical density OD_600_ 0.2-0.3 and 1.6, respectively, measured with NovaspecII Visible Spectrophotometer, Pharmacia Biotech). For amino acids auxotrophy tests cells were plated on 1.5% (wt/vol) agar with Spizizen minimal salts (SM; [150] supplemented with 0.5% (wt/vol) glucose, 0.1% (wt/vol) glutamate and 0.5% or 0.004% (wt/vol) Casamino Acids. Standard protocols were used for transformation of *E. coli* and *B. subtilis* competent cells [150].

Sporulation was analyzed in supplemented Difco Sporulation medium (Difco) [151]. To determine the level of ppGpp, cells were grown in the defined SM medium. When required for selection, antibiotics were added at following concentrations: 100 *μ*g per ml of ampicillin, 100 *μ*g per ml of spectinomycin, 0.5 *μ*g per ml of erythromycin, 3 µg per ml of phleomycin, 5 µg per ml of kanamycin, and 5 *μ*g per ml of chloramphenicol. IPTG (isopropyl-β-D-1-thiogalactopyranoside) inducer was added to cells at concentrations indicated in the main text.

### Strains and plasmid construction

*E. coli* TG1 strain was used for plasmids construction. All *B. subtilis* constructions were performed at the basis of BSB1 strain. The used oligonucleotides are listed in S6 Table. To construct the system for stable Rho expression, *rho* open reading frame was fused by PCR to the ribosome-binding site and spacer sequence of *B. subtilis tagD* gene using BSB1 chromosome as a template and oligonucleotides eb424 and eb458. The amplified fragment was cloned downstream P*_veg_* promoter at pDG1730 plasmid using the blunted NheI and EagI sites. The resulting plasmid was transformed into *B. subtilis* BSB1 cells with selection for spectinomycin-resistance leading to the integration of the P*veg-rho* expression unit at chromosomal *amyE* locus by double crossover.

To construct a similar system expressing the tagged Rho, *rho-SPA* DNA fragment was amplified from the chromosome of BRL415 strain [8] using oligonucleotides eb458 and op148-R, digested by EagI endonuclease and cloned at pDG1730 between EagI and the filled-in NheI sites. The P*veg-rho*-SPA expression unit was inserted into the BSB1 chromosome as above.

To overexpress Rho in the strain BRL1250 containing P*_hy-spanc_-kinC* fusion (Spec^R^), P*_veg_-rho* expression unit was amplified from the chromosome of BRL802 strain using the oligonucleotides opv1730-B and veb596, digested by BamHI endonuclease and cloned between BamHI and blunted EcoRI sites of pSWEET vector. The resulting plasmid was used to reinsert the P_veg_-rho expression unit at the *amyE* locus of BSB1 strain as above with selection for chloramphenicol-resistance. The resulting strain BRL1248 was controlled for sporulation- and competence-negative phenotypes associated with Rho overexpression. Finally, Pveg-rho fusion was transformed into BRL1250 cells with selection for chloramphenicol-resistance.

The *B. subtilis* partial loss-of-function mutants *guaB S121F* and *guaB T139I* were constructed as follows. The corresponding single-nucleotide mutations *c362t* and *c416t* (coordinates starting from the *guaB* +1 nucleotide) were introduced into the *guaB* gene by the two-step site-directed mutagenesis. First, the DNA fragments were PCR-amplified using the complementary mutagenic oligonucleotides veb852, veb853 (for *c362t*), and veb855, veb856 (for *c416t*) in pairs with correspondent primers veb857, veb858 (for *c362t*), and veb857, veb859 (for *c416t*). Next, the respective fragments were joined by PCR using the primers veb857 and veb858 (for *c362t*) and veb857 and veb859 (for *c416t*) and cloned at the thermo-sensitive shuttle plasmid pMAD [152] between SalI and EcoRI sites. The resulting plasmids were transformed in BSB1 cells with the selection for erythromycin-resistance at non-permissive 37°C. In this way, the plasmids integrated at the chromosomal *guaB* locus by single crossover leading to *guaB* duplication. The selected clones were propagated without selection at permissive 30°C to induce plasmid replication and its segregation from the chromosome due to the recombination between the flanking *guaB* copies. The erythromycin-sensitive clones which lost the plasmid were tested for the presence of *guaB* mutations by PCR using common primer veb857 and the oligonuclotides veb851 and veb854, specific for *c362t* and *c416t* mutations, respectively. The selected *guaB* mutants were controlled by sequencing.

### Luciferase assay

Analysis of promoters’ activity using luciferase fusions was performed as described previously with minor modifications [52]. Cells were grown in LB medium to mid-exponential phase (optical density OD_600_ 0,4-0,5 with NovaspecII Visible Spectrophotometer, Pharmacia Biotech), after which cultures were centrifuged and resuspended to OD 1.0 in fresh DS media, to follow expression of *spo0A-luc* and s*poIIA-luc* fusions during sporulation, or in competence-inducing MM media, to analyze *comK-luc* activity during competence development. Upon OD verification, these pre-cultures were next diluted in respective media to an OD_600_ 0.025. The starter cultures were distributed by 200µl in a 96-well black plate (Corning, USA) and Xenolight D-luciferin K-salt (Perkin, USA) was added to each well to final concentration 1.5 mg/mL. The cultures were incubated at 37°C with agitation and analyzed in Synergy 2 Multi-mode microplate reader (BioTek Instruments). Relative Luminescence Units (RLU) and OD_600_ were measured at 5 min intervals. Each fusion-containing strain was analyzed at least three times. Each experiment included four independent cultures of each strain.

### Epifluorescence microscopy, image processing and cell measurements

Cultures of *B. subtilis* were performed as described above. Cultures were sampled during exponential growth (OD_600_ 0.2) and stationary phase (OD_600_ 1.3), and cells were mixed with Nile Red (10 µg/ml final concentration) before mounting on a 2% agarose pad and topped with a coverslip. Bacteria were imaged with an inverted microscope (Nikon Ti-E), controlled by the MetaMorph software package (v 7.8; Molecular Devices, LLC), equipped with a 100× oil immersion phase objective. Epifluorescence images were recorded on phase-contrast and fluorescence channels (ex. 562 ± 40/em. 641 ± 75 nm filters) with an ORCA-R2 camera (Hamamatsu), and 100 ms exposure time. The post-acquisition treatment of the images was done with the Fiji software [153, 154]. The mean cell lengths were determined with the ChainTracer plugin of the Fiji software [155] on two independent experiments with N >140 (N_avg_ = 290).

### Sporulation assay

For sporulation assay, cells were diluted in LB in a way to obtain the exponentially growing cultures after over-night incubation at 28°C. The pre-cultures were diluted in pre-warmed liquid DS medium at OD600 0.025 and incubated at 37°C for 20 or 24 hours. To determine quantity of the spores, half of a culture was heated at 75°C for 15 min and cells from heated and non-heated samples were platted in sequential ten-fold dilutions at LB-agar plates. Colonies were counted after 36 h of incubation at 37°C, and the percentage of spores was calculated as the ratio of colonies forming units in heated and unheated samples. In the sporulation experiments employing the IPTG-inducible systems for *kinA* or *kinC* expression, cells were let to sporulate in the presence of IPTG at concentrations indicated in the text.

Each experiment included three independent isogenic cultures. Four independent experiments were performed to establish sporulation efficiency of each strain.

### Genetic competence assay

To establish kinetics of competence development using a two-step transformation procedure [150]. *B. subtilis* cells were grown in a rich defined medium SpC to stationary phase (OD_600_ 1.5) and diluted 7-fold in a competence-inducing medium SpII; at 30-min intervals, culture samples (0.25 ml) were mixed with *B. subtilis* BSF4217 genomic DNA (100 ng) or pIL253 plasmid DNA (500 ng), incubated for 30 min at 37°C and platted at LB plates containing erythromycin. Plates were incubated at 37°C for 18 h before colonies counting.

### Western blotting

The crude cell extracts were prepared using Vibracell 72408 sonicator (Bioblock scientific). Bradford assay was used to determine total protein concentration in each extract. Equal amounts of total proteins were separated by SDS-PAGE (10% polyacrylamide). After the run, proteins were transferred to Hybond PVDF membrane (GE Healthcare Amersham, Germany), and the transfer quality was evaluated by staining the membrane with Ponceau S (Sigma-Aldrich). The SPA-tagged Rho protein was visualized by hybridization with the primary mouse ANTI-FLAG M2 monoclonal antibodies (Sigma-Aldrich; dilution 1:5,000) and the secondary goat peroxidise-coupled anti-mouse IgG antibodies A2304 (Sigma-Aldrich; dilution 1:20,000). The control Mbl protein was visualized using primary rabbit anti-Mbl antibodies (dilution 1:10,000) and the secondary goat peroxidase-coupled anti-rabbit IgG antibodies A0545 (Sigma-Aldrich; dilution 1:10,000). Three independent experiments were performed, and a representative result is shown in Fig 1B

### ppGpp determination

To determine intracellular ppGpp level, *B. subtilis* cells were grown in the defined MS medium supplemented with 0.5% (wt/vol) glucose, 0.1% (wt/vol) glutamate and 0.5% (wt/vol) Casamino Acids to optical densities OD_600_ 0.5 (for argenine hydroxamate treatment analysis) or OD_600_ 1.5 (for the stationary phase analysis).

Bacterial cultures in triplicates (20 ml each) were rapidly centrifuged at 4°C and cellular pellets were frozen in liquid nitrogen. All extraction steps were performed on ice. Cellular pellets were deproteinized by addition of an equal volume of 6% perchloric acid (PCA) and incubation on ice for 10 min with two rounds of vortex-mixing for 20 s. Acid cell extracts were centrifuged at 13,000 rpm for 10 min at 4°C. The resulting supernatants were supplemented with an equal volume of bi-distilled water, vortex-mixed for 60 s, and neutralized by addition of 2 M Na_2_CO_3_. After filtration (3kDa cut off), extracts were injected onto a C18 Supelco 5 µm (250 × 4.6 mm) column (Sigma) at 45°C. The mobile phase was delivered using the stepwise gradient of buffer A (10 mM tetrabutylammonium hydroxide, 10 mM KH2PO4 and 0.25% MeOH; adjusted with 1M HCl to pH 6.9) and buffer B (5.6 mM tetrabutylammonium hydroxide, 50 mM KH2PO4 and 30% MeOH; adjusted with 1 M NaOH to pH 7.0) at a flow-rate of 1 ml/min and elution program: from 60%A + 40%B at 0 min to 40%A+60%B at 30 min and 40%A+60%B at 60 min.

Detection was done with a diode array detector (PDA). The LC Solution workstation chromatography manager was used to pilot the HPLC instrument and to process the data. Products were monitored spectrophotometrically at 254 nm, and quantified by integration of the peak absorbance area, employing a calibration curve established with various nucleoside standards. ppGpp standard was purchased from Jena Bioscience GmbH (Germany). Finally, a correction coefficient was applied to correct raw data for minor differences in the densities of bacterial cultures.

### Transcriptome profiling by RNA sequencing

RNA was extracted from independent cultures of *B. subtilis* BsB1 WT, Δ*rho* and Rho^+^ strains grown in LB medium at 37°C under vigorous agitation up to mid exponential or early stationary phase of growth (OD600 ∼0.5 and ∼2.0, respectively). Experiments were performed in duplicates for WT and mid-exponential samples and triplicates for early stationary phase of Δ*rho* and Rho^+^.

RNA preparation and DNase treatment were done as described [6]. Quality and quantity of RNA samples were analyzed on Bioanalyzer (Agilent, CA). The Next Generation Sequencing (NGS) Core Facility (Institute of Integrative Biology of the Cell, Gif-sur-Yvette, France; https://www.i2bc.paris-saclay.fr/sequencing/ng-sequencing/addon-ng-sequencing) prepared the RNAseq libraries with ScriptSeq protocol using RiboZero for rRNA-depletion (Illumina, San Diego, California) and generated strand-specific paired-end reads of 40 bp on an Illumina NextSeq platform (NextSeq 500/550 High Output Kit v2).

Reads were trimmed to remove adapters and low-quality ends using Cutadapt (v1.15, DOI:10.14806/ej.17.1.200) and Sickle (v1.33, options: -t sanger -x -n -q 20 -l 20) and mapped onto AL009126.3 reference genome assembly using Bowtie2 (v2.3.5.1; options “-N 1 -L 16 R 4”, [156]. Counts of the number of read pairs (fragments) overlapping the sense and antisense strand of each transcribed region (AL009126.3-annotated genes and S-segments from [6] were obtained with Htseq-count (v0.11.0; options “-m union –nonunique=all”; 156 Anders *et al*., 2015).

Since Δ*rho*, WT and Rho^+^ exhibited different levels of pervasive transcription leading to global changes in low expression values and antisense signal; we selected a subset of well-expressed genes whose sense signal is in principle less impacted and thus most relevant for sequencing depth normalization. To this end, we selected the 728 AL009126.3-annotated genes satisfying, for all 4 WT samples, log2(fpkm_raw+5)>7, where fpkm_raw refers to the fpkm (fragments per kilobase of transcript per million mapped fragments) value obtained when library size is simply estimated of as the sum of counts. Differential gene expression analysis between conditions and strains, including sequencing depth normalization, was then conducted with R library “DESeq2” (v1.32.0; [158]). DESeq2 p-values for each pairwise comparison and each strand were converted into q-values using R library “fdrtool” (v1.2.17; [159]. Genes were called differentially expressed between strains or conditions when the estimated q-value ≤ 0.05 and |log2FC| exceeded the cut-off specified in the text (0.5 or 1) for the considered strand (sense or antisense). Data was deposited in GEO (accession number GSE195579).

Graphical representations of the expression level of a gene in a given strain and condition used the geometrical mean of log2(fpkm+5) values, where FPKM was computed with the DESeq2-estimated library size factors multiplied by the median of sample count sums. To allow interactive exploration of the sense and antisense signal along the genome with bp-resolution, we also implemented in Genoscapist [76] the representation of a new data type corresponding to RNAseq coverage. For this purpose, count values are extracted with “bedtools genomecov” (version 2.27.0, 156.; [160]) and represented as a step function with breakpoints corresponding to extremities of mapped read pairs along the genome sequence. To make these counts comparable between different samples and with gene-level expression values, coverage counts are converted to fpkm, using the formula fpkm_cov(t) = cov(t)*(10^3^/F)*(10^6^/L), where cov(t) is the coverage count for genome position t, L is the library size used to compute gene-level FPKM, and F is the average fragment length (from 178 bp to 200 bp across samples) obtained from the distance between extremities of inward oriented read pairs returned by “samtools stats” (version 1.10, 157.; [161]). The bp-level signal, displayed as log2(fpkm_cov(t)+5), can be accessed via the website http://genoscapist.migale.inrae.fr/seb_rho/.

## Acknowledgments

This work has benefited from the facilities and expertise of the high throughput sequencing core facility of I2BC (Research Center of GIF – http://www.i2bc.paris-saclay.fr/.). We are grateful to the INRAE MIGALE bioinformatics facility (doi: 10.15454/1.5572390655343293E12) for providing computing and storage resources.

## Supporting Information

**S1 Fig. Graphical representation of the sporulation-proficient suppressor mutations in Rho from *B. subtilis* Rho*^+^* strain.** The primary sequence of *B. subtilis* Rho subunit is shown (NP_391589.2). The major motifs (as in D’Heygère *et al*., 2015) are boxed and highlighted in grey. The identified amino acid substitutions are marked in red.

**S2 Fig. *B. subtilis* Rho^+^ exhibits competence–negative phenotype.** Transformability of *B. subtilis* WT (blue lines) and Rho^+^ (red lines) strains by the plasmid pIL253 (closed circles) and homologous genomic DNA (open circles). Competence induction and transformation were performed as described in Materials and Methods and Fig 3B. The experiment included three biological replicas of each strain and was reproduced twice. The results of a representative experiment are presented. The data are independent from Fig 3B.

**S3 Fig. Rho^+^ strain differs from *spo0A* mutant in the activation of *comK*.** Kinetics of luciferase expression in *B. subtilis* WT (blue lines), Rho^+^ (red lines) and *spo0A* (gray lines) mutant cells bearing the P*_comK_-luc* transcription fusion and grown in competence-inducing medium as described in Materials and Methods. For each strain, plotted are the mean values of luminescence readings corrected for OD from four independent cultures analyzed simultaneously. The double-lined curves depict characteristic growth kinetics of cells measured by OD_600_. The experiment was reproduced three times and is independent from Fig 2C. The data from a representative experiment are presented.

**S4 Fig. CodY inactivation does not restore *comK* expression in Rho^+^ cells.** Kinetics of luciferase expression from the P*_comK_-luc* transcription fusion in *B. subtilis* WT (blue open squares), Rho^+^ (red open circles) cells and their respective *codY* mutants (fill-in blue and red symbols) grown in competence-inducing medium. For each strain, plotted are the mean values of luminescence readings corrected for OD from four independent cultures analyzed simultaneously. Characteristic growth kinetics of WT and Rho^+^ cells and of their *codY* derivatives are depicted by double- and single-lined curves, respectively. Presented are the results of two independent experiments performed using freshly prepared media.

**S5 Fig. Genome wide effects of Rho over-production on the *B. subtilis* transcriptome during exponential growth and stationary phase in rich medium.** Transcriptome changes in the antisense (A and B) and sense (C and D) strands during exponential growth (A and C) and stationary phase (B and D), respectively. Each point represents one of the 4,292 AL009126.3-annotated genes. Coordinates on x- and y-axes correspond to the normalized expression level (average of log2(fpkm+5) over biological replicates) measured with RNAseq in *B. subtilis* WT and Rho^+^, respectively. Background colors of the points indicate TRs whose transcription level is strongly up-regulated (yellow) or down-regulated (blue) in the Rho^+^ vs. WT comparison made in *B. subtilis* by RNAseq.

**S6 Fig. Differential expression of the ComK, CodY and stringent response regulons in *B. subtilis* Δ*rho* strain.** Expression changes for each gene from the ComK regulon (**A, B**), CodY and the stringent response regulons (**C, D**) by comparing *B. subtilis* Δ*rho* and WT strains under exponential growth (A and C) and stationary phase (B and D) conditions. Each point (red for the ComK regulon, blue for the CodY regulon, yellow for the stringent response regulon, and gray for genes outside of the mentioned regulons) represents one of the 4,292 AL009126.3-annotated genes.

**S7 Fig. Lowering GTP levels does not rescue thermo-sensitive phenotype of Rho^+^ strain.** *B. subtilis* WT, Rho^+^ cells and their respective *guaB* S121F and *guaB* T139I mutants were grown in LB medium at 37°C to mid exponential phase (OD_600_ 0.5), spotted in serial dilutions on LB agar plates and incubated at 37°C and 55°C for 18 hours. The experiment was reproduced at least three times and the representative results are shown.

**S8 Fig. Thermo-sensitivity of Rho^+^ strain is not due to a low level of *comGA* expression, as *comGA* mutant resists high temperature.** *B. subtilis* WT, its isogenic *comK* and *comGA* mutants and Rho^+^ cells growing exponentially (OD_600_ 0.5) in LB medium were streaked on LB agar plates and incubated at 37°C and 55°C for 18 hours before imagining. One representative experiment out of three conducted is shown.

**S1 Table Sporulation proficient and thermoresistant suppressors of Rho^+^ strain.**

**S2 Table Differential expression analysis of Rho^+^ and Δ*rho* vs. WT *B. subtilis* BsB1 cells.**

**S3 Table Comparison between sets of DE genes and *Subti*Wiki regulons.**

**S4 Table. Strains and plasmids used in this study.**

**S5 Table. Oligonucleotides used for strains construction.**

