## Supplemental figures for "Termination factor Rho mediates transcriptional reprogramming of *Bacillus subtilis* stationary phase"

RNA binding domain

MKDVSISSLENMKLKELYELARHYKISYYSKLTKKELIFAILKANAEQEDLLFMEGVLEIIQSEGFGFLRPINYSPSSEDI

stop

YISASQIRRFDLRNGDKVSGKVRPPKENERYYGLLHVEAVNGDDPESAKERVHFPALTPLYPDRQMVLETKPNFLST

T

RIMDMMAPVGFGQRGLIVAPPKAGKTMLLKEIANSITANQPEAELIVLLIDERPEEVTDIERSVAGDVVSSTFDEVP

Walker A

H RR

ENHIKVAELVLERAMRLVEHKKDVIILMDSITRLARAYNLVIPPSGRTLSGGIDPAAFHRPKRFFGAARNIEEGGSLTIL

Walker B Q-loop

R

ATALVDTGSRMDDVIYEEFKGTGNMELHLDRSLAERRIFPAIDIRRSGTRKEELLVPKEHLDRLWSIRKTMSDSPDFA

R-loop Arg finger

EKFMRKMKKTKTNQEFFDILNQEWKQANLSSARR

S1 Fig

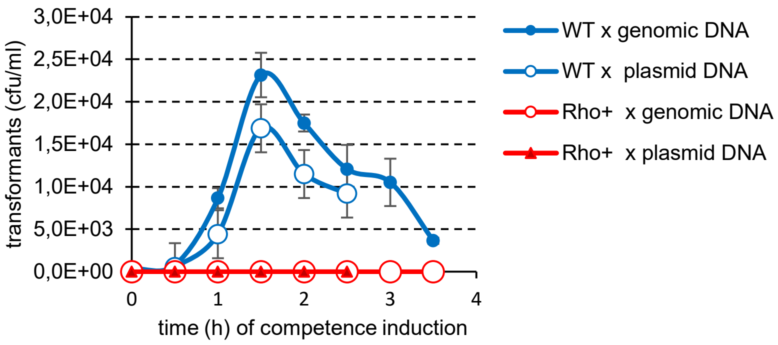

3,0E+04
2,5E+04
2,0E+04
1,5E+04
1,0E+04
5,0E+03
0,0E+00
transformants (cfu/ml)
0
1
2
3
4
time (h) of competence induction
WT x genomic DNA
WT x plasmid DNA
Rho+ x genomic DNA
Rho+ x plasmid DNA


S2 Fig

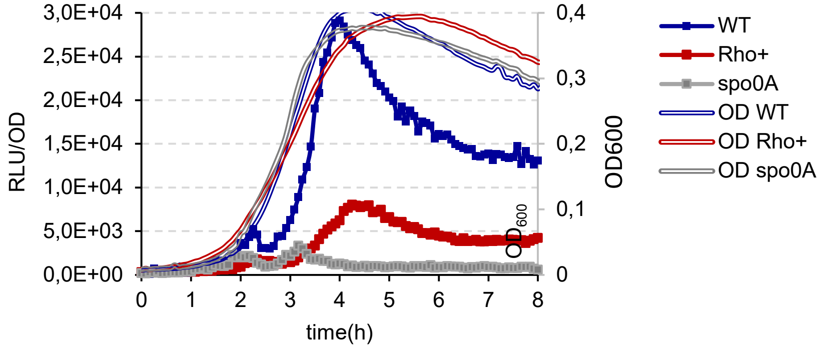
3,0E+04
2,5E+04
2,0E+04
1,5E+04
1,0E+04
5,0E+03
0,0E+00
RLU/OD
0,4
0,3
0,2
0,1
0
OD600
$OD_{600}$
0 1 2 3 4 5 6 7 8
time(h)
WT
Rho+
spo0A
OD WT
OD Rho+
OD spo0A

S3 Fig

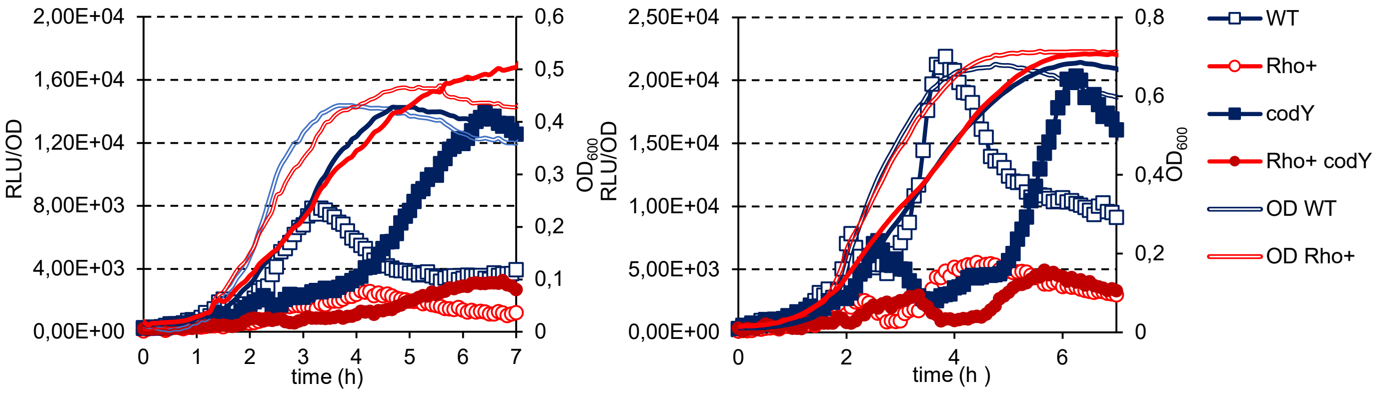

2,00E+04
1,60E+04
1,20E+04
8,00E+03
4,00E+03
0,00E+00
RLU/OD
0,6
0,5
0,4
0,3
0,2
0,1
0
OD600
0 1 2 3 4 5 6 7
time (h)
2,50E+04
2,00E+04
1,50E+04
1,00E+04
5,00E+03
0,00E+00
RLU/OD
0,8
0,6
0,4
0,2
0
OD600
0 2 4 6
time (h )
WT
Rho+
codY
Rho+ codY
OD WT
OD Rho+


S4 Fig

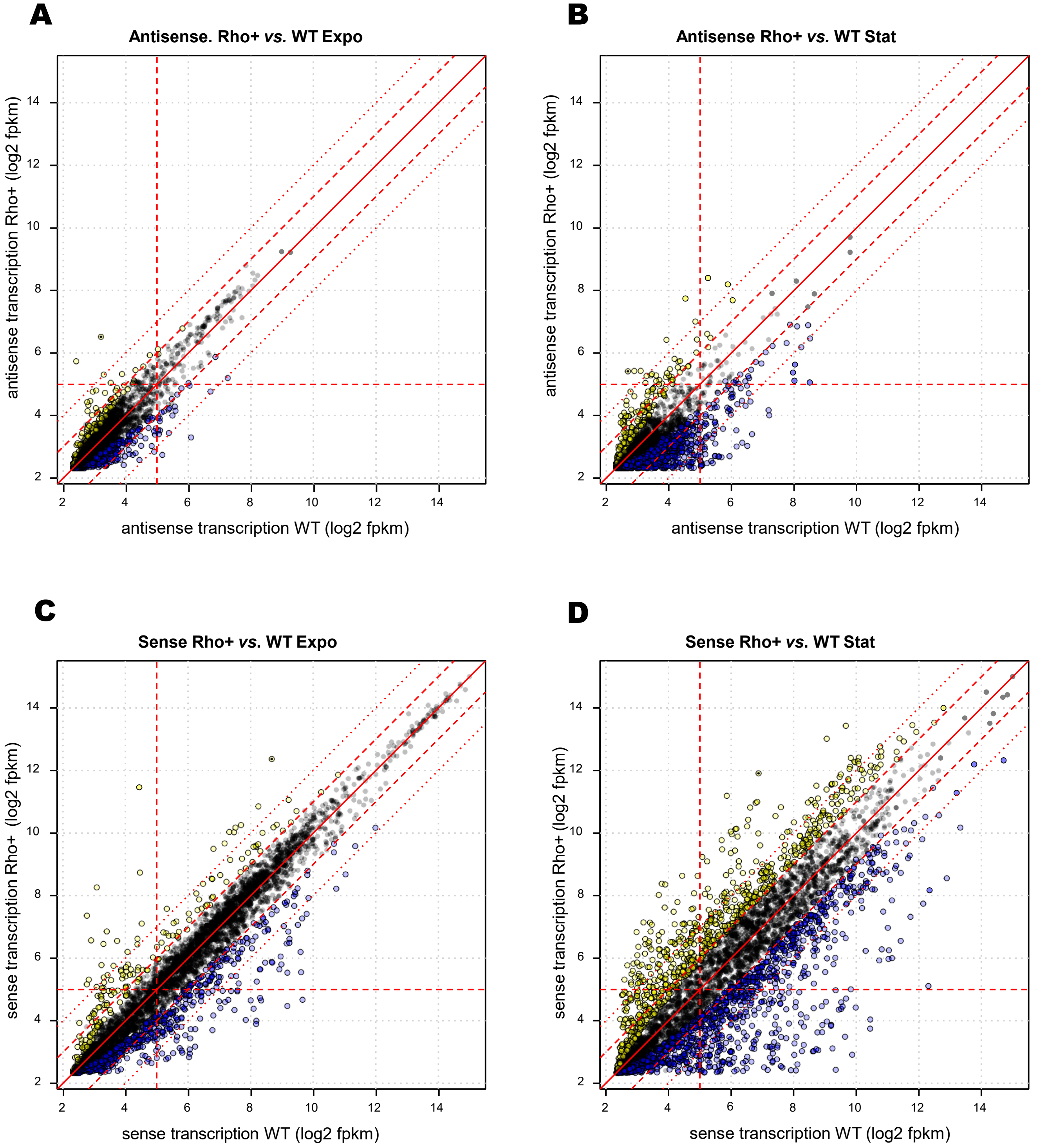
A
Antisense. Rho+ vs. WT Expo
antisense transcription Rho+ (log2 fpkm)
antisense transcription WT (log2 fpkm)
B
Antisense Rho+ vs. WT Stat
antisense transcription Rho+ (log2 fpkm)
antisense transcription WT (log2 fpkm)
C
Sense Rho+ vs. WT Expo
sense transcription Rho+ (log2 fpkm)
sense transcription WT (log2 fpkm)
D
Sense Rho+ vs. WT Stat
sense transcription Rho+ (log2 fpkm)
sense transcription WT (log2 fpkm)
2
4
6
8
10
12
14

S5 Fig

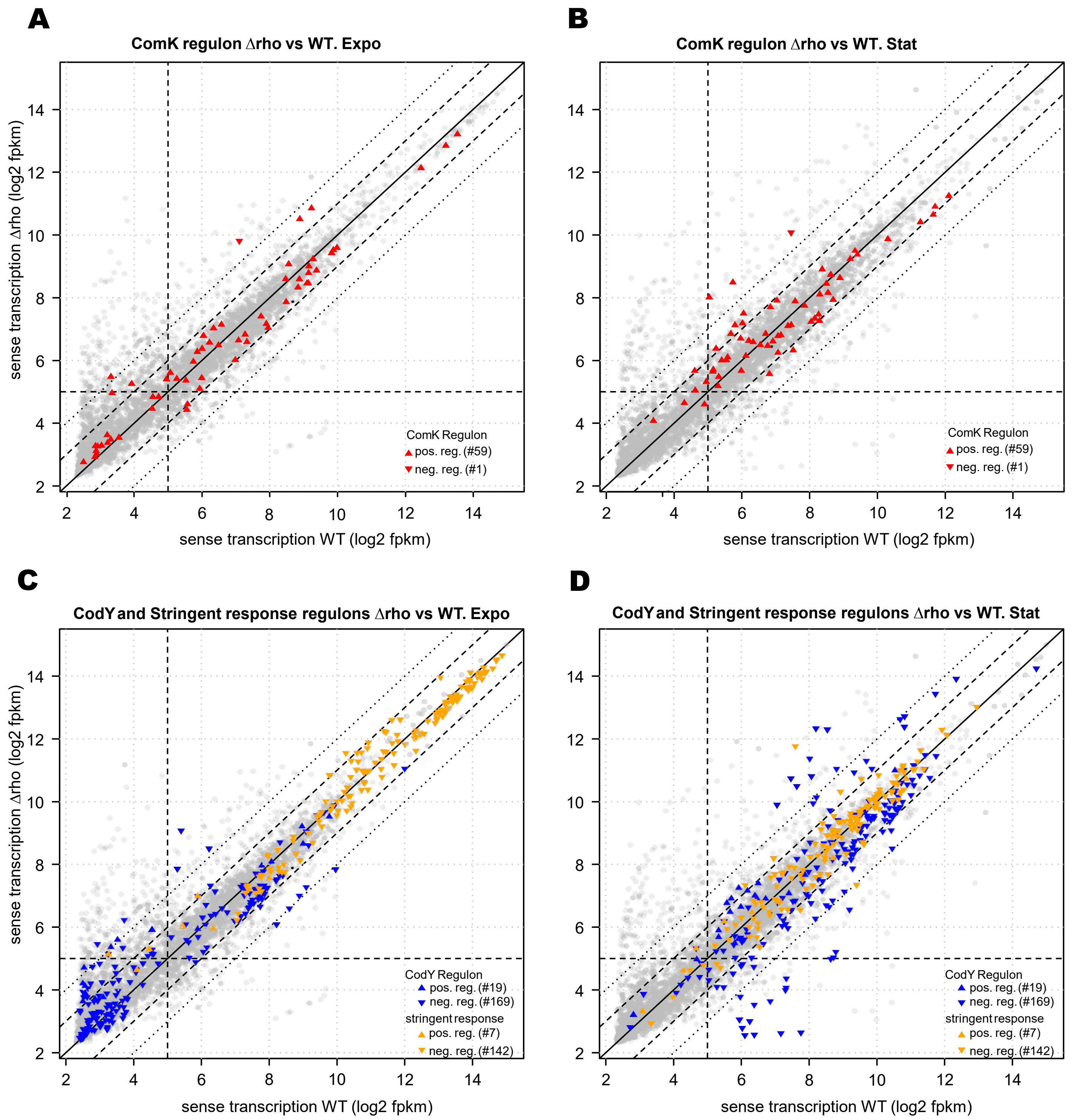
A
ComK regulon Δrho vs WT. Expo
B
ComK regulon Δrho vs WT. Stat
C
CodY and Stringent response regulons Δrho vs WT. Expo
D
CodY and Stringent response regulons Δrho vs WT. Stat
sense transcription Δrho (log2 fpkm)
sense transcription WT (log2 fpkm)
2
4
6
8
10
12
14
ComK Regulon
pos. reg. (#59)
neg. reg. (#1)
CodY Regulon
pos. reg. (#19)
neg. reg. (#169)
stringent response
pos. reg. (#7)
neg. reg. (#142)

S6 Fig

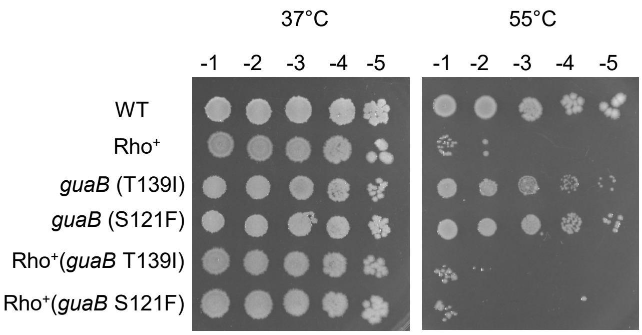

37°C
55°C
-1 -2 -3 -4 -5
-1 -2 -3 -4 -5
WT
Rho+
*guaB* (T139I)
*guaB* (S121F)
Rho+(*guaB* T139I)
Rho+(*guaB* S121F)


S7 Fig

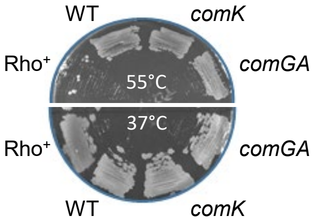


S8 Fig
