## Supplemental Table 1 for "Termination factor Rho mediates transcriptional reprogramming of *Bacillus subtilis* stationary phase"

### S1 Table

#### Sporulartion proficient suppressors

| | sporulation efficiency | | | propeties of the mutated ectopic rho expression unit | | DNA mutation(****): | Protein change |
|---|---|---|---|---|---|---|---|
| | initial isolate | transfered to RM (*Δ rho*) | transfered to WT | *Δ rho* complementation | relationship to the rho copy at locus | nucleotide substitution | |
| Rho+ supp1 | 6,54E-01 | 8,57E+00 | 1,52E+00 | no(*) | recesive (**) | c436t | Gln146Stop |
| Rho+ supp2 | 3,49E-01 | 8,53E-01 | 8,44E-01 | yes | recesive | c1064g | Pro335Arg |
| Rho+ supp3 | 2,80E+00 | 5,26E+00 | 4,02E+00 | no | negatively dominant (***) | g856a | Gly286Arg |
| Rho+ supp4 | 4,42E-01 | 9,85E+00 | 6,93E-01 | no | recesive | c436t | Gln146Stop |
| Rho+ supp5 | 6,81E-01 | 1,09E+00 | 1,19E+00 | yes | recesive | a820c | Asn274His |
| Rho+ supp6 | 4,18E+00 | 3,86E+00 | 5,17E+00 | no | negatively dominant | g859a | Gly287Arg |
| Rho+ supp7 | 4,97E+00 | 4,74E+00 | 5,34E+00 | no | negatively dominant | g529a | Ala177Thr |
| Rho+ supp8 | 1,12E+00 | 1,26E+00 | 1,40E+00 | yes | recesive | a820c | Asn274His |
| **controls** | | | | | | | |
| wt | 1,00E+00 | | | | | | |
| RM | 1,02E+01 | | | | | | |
| Rho+ | 2,87E-04 | | | | | | |

(*) when transfered to RM cells, mutated Rho expressed ectopicaly from Pveg does not reduce excessive sporulation characteristic to RM.
(**) when transfered to WT cells, mutated Rho expressed ectopicaly from Pveg does not modify sporulation efficiency significantly.
(***) when transfered to WT cells, mutated Rho expressed ectopicaly from Pveg increases sporulation efficiency above the WT level; probably due to "poisoning" of Rho expressed at locus from natural promoter.
(****) Position of DNA mutations from the +1 nucleotide of *rho* .

#### Thermoresistant suppressors obtained at 55°C

| | DNA mutation(****): nucleotide deletion or insertion | Protein change |
|---|---|---|
| Rho+ suppTR1 | a inserted at 1207 | Thr403Asn -> 39 novel C-terminal amino acids |
| Rho+ suppTR2 | a inserted at 100 | Thr36Arg->Stop at aa64 |
| Rho+ suppTR3 | deletion g932 | Ser311Thr->Stop at aa312 |
| Rho+ suppTR4 | deletion g1200 | Met400Ile-> 34 novel C-terminal amino acids |

(****) Position of DNA mutations from the +1 nucleotide of *rho* .
