## Supplemental Table 4 for "Termination factor Rho mediates transcriptional reprogramming of *Bacillus subtilis* stationary phase"

**S4 Table. Strains and plasmids used in this study.**

| Strains | Genotype (resistance) | Source or reference |
|---|---|---|
| BSB1 | *B. subtilis* 168 *trp*$^{+}$ | Nicolas et al., 2012 |
| BRL1 | BSB1 *Δrho::phleo* | Bidnenko et al., 2017 |
| BRL802 (Rho$^{+}$ spec) | BSB1 *amyE::Pveg-rho* (Sp$^{R}$) | This study |
| BRL415 | BSB1 *rho-SPA* (Em$^{R}$) | Bidnenko et al., 2017 |
| BRL796 | BSB1 *amyE::Pveg-rho-SPA* (Sp$^{R}$) | This study |
| BRL116 | BSB1 P*spo0A-luc* (Cm$^{R}$) | Bidnenko et al., 2017 |
| BRL831 | Rho$^{+}$ P*spo0A-luc* (Cm$^{R}$ Sp$^{R}$) | This study |
| BKK00980 | *B. subtilis* 168 *trpC2 spo0H::kan* (Km$^{R}$) | Koo et al., 2017 |
| BRL896 | BSB1 *spo0H::kan* P*spo0A-luc* | This study |
| BRL111 | BSB1 P*spoIIA-luc* (Cm$^{R}$) | Bidnenko et al., 2017 |
| BRL809 | Rho$^{+}$ P*spoIIA-luc* (Cm$^{R}$) | This study |
| MF1913 | *B. subtilis* PY79 $P_{hyspanc}$-kinA (CmR) | Fujita and Losick, 2005 |
| BRL1240 | BSB1 $P_{hyspanc}$-kinA (Cm$^{R}$) | This study |
| BRL1241 | Rho+ $P_{hyspanc}$-kinA (CmR Sp$^{R}$) | This study |
| RL3606 | *B. subtilis* PY79 kinCΩ$P_{hyspanc}$-kinC (Sp$^{R}$) | Fujita et al., 2005 |
| BRL1244 | BSB1 kinCΩ$P_{hyspanc}$-kinC (Sp$^{R}$) | This study |
| BRL1248 (Rho$^{+}$ cat) | BSB1 *amyE::Pveg-rho* (Cm$^{R}$) | This study |
| BRL1250 | Rho$^{+}$ kinCΩ$P_{hyspanc}$-kinC (Sp$^{R}$Cm$^{R}$) | This study |
| BD4773 | *B. subtilis* P*comK-luc* (Cm$^{R}$) | Mirouze et al., 2012 |
| BRL115 | BSB1 P*comK-luc* (Cm$^{R}$) | This study |
| BRL829 | Rho$^{+}$ P*comK-luc* (Sp$^{R}$ Cm$^{R}$) | This study |
| BKK14240 | *B. subtilis* 168 *trpC2 rok::kan* (Km$^{R}$) | Koo et al., 2017 |
| BRL1273 | BSB1 *rok::kan* (Km$^{R}$) | This study |
| BRL1274 | Rho$^{+}$ *rok::kan* (Km$^{R}$ Sp$^{R}$) | This study |
| BRL1303 | BSB1 P*comK-luc rok::kan* (Cm$^{R}$ Km$^{R}$) | This study |
| BRL1304 | Rho$^{+}$ P*comK-luc rok::kan* (Cm$^{R}$ Km$^{R}$ Sp$^{R}$) | This study |
| BKK16170 | *B. subtilis* 168 *trpC2 codY::kan* (Km$^{R}$) | Koo et al., 2017 |
| BRL1126 | BSB1 P*comK-luc codY::kan* (Cm$^{R}$ Km$^{R}$) | This study |
| BRL1133 | Rho$^{+}$ P*comK-luc codY::kan* (Cm$^{R}$ Km$^{R}$ Sp$^{R}$) | This study |
| BRL1267 | BSB1 *guaBS121F* | This study |
| BRL1268 | BSB1 *guaBT139I* | This study |
| BRL1271 | Rho$^{+}$ *guaBS121F* (Sp$^{R}$) | This study |
| BRL1272 | Rho$^{+}$ *guaBT139I* (Sp$^{R}$) | This study |
| CCB1050 *(p)ppGpp$^{0}$* | *B. subtilis* W168 *yjbM::spc ywaC::kan relA::ery* (Em$^{R}$ Km$^{R}$ Sp$^{R}$) | Trinquier et al., 2019; Kriel et al., 2012 |
| **Plasmids** | | |
| pMAD | Erm$^{R}$, Amp$^{R}$ | Arnaud et al., 2004 |
| pDG1730 | Sp$^{R}$, Amp$^{R}$ | Guérout-Fleury et al., 1996 |
| pSWEET | Cm$^{R}$, Amp$^{R}$ | Bhavsar et al., 2001 |
| pDG1730-rho | Pveg-rho at pDG1730 ; Sp$^{R}$, Amp$^{R}$; used to construct BRL802 | This study |

| | | |
|---|---|---|
| pDG1730-rho-SPA | Pveg-rho-SPA at pDG1730 ; $Sp^R$, $Amp^R$; used to construct BRL796 | This study |
| pSWEET-Pveg-rho | Pveg-rho at pSWEET ; $Cm^R$, $Amp^R$; used to construct BRL1248 | This study |
| pMAD*guaBS121F* | $Erm^R$, $Amp^R$; used to construct BRL1267 | This study |
| pMAD*guaBS139I* | $Erm^R$, $Amp^R$; used to construct BRL1268 | This study |

1. Arnaud M, Chastanet A, Débarbouillé M. New vector for efficient allelic replacement in naturally nontransformable, low-GC-content, Gram-positive bacteria. Appl Environ Microbiol. 2004; 70: 6887-6891. doi:10.1128/AEM.70.11.6887-6891.2004
2. Guérout-Fleury A-M, Frandsen N, Stragier P. Plasmids for ectopic integration in Bacillus subtilis. Gene 1996; 180(1-2): 57-61. https://doi.org/10.1016/S0378-1119(96)00404-0
3. Bhavsar, A. P., Zhao, X., Brown, E. D. Development and characterization of a xylose-dependent system for expression of cloned genes in Bacillus subtilis: conditional complementation of a teichoic acid mutant. Appl Environ Microbiol. 2001; 67(1) : 403–410. https://doi.org/10.1128/AEM.67.1.403-410.2001
