## Supplemental Table 5 for "Termination factor Rho mediates transcriptional reprogramming of *Bacillus subtilis* stationary phase"

**S5 Table. Oligonucleotides used for strains construction**

| Oligonucleotide | Sequence (5'-> 3') |
|---|---|
| eb458 (*) | GCGCTTAATTAACAT**TAGGAAGGAGCG**TTTCTTTAATGAAAGACGTATCTATTTCC |
| eb424 (**) | CG*GGATCC*TTACCTTCTTGCAGATGATAG |
| op148-R | GTA*CGTACG*ATCTTTCAGCCG |
| opv1730-B | TTC*GGATCC*TGTATTACTATTC |
| veb596 | GC*GTCGAC*TTACCTTCTTGCAGATGATAG |
| veb851(***) | ATGGGGAAATACAGAATTT**T** |
| veb852 (****) | ATACAGAATTT**T**CGGTGTTCCG |
| veb853 | AACACCG**A**AAATTCTGTATTTC |
| veb854 (***) | AAGCTTGTTGGAATTATTA**T** |
| veb855 | GGAATTATTA**T**AAACCGTGACC |
| veb856 | GGTCACGGTTT**A**TAATAATTCC |
| veb857 | GTT*GAATTC*AACGGAGTAAAATAG |
| veb858 | CTT*GTCGAC*TTCAGCTGTTG |
| veb859 | TCA*GTCGAC*AATCATAAATTGC |

* Ribosome binding site and spacer sequence of tagD gene are bolded and underlined, respectively.

** Here and thereafter the sites of endonuclease are in italics.

*** The 3'-terminal nucleotides matching the mutations are bolded.

**** Here and thereafter the mutated nucleotides are bolded and underlined.
